# Foundation model of neural activity predicts response to new stimulus types and anatomy

**DOI:** 10.1101/2023.03.21.533548

**Authors:** Eric Y. Wang, Paul G. Fahey, Zhuokun Ding, Stelios Papadopoulos, Kayla Ponder, Marissa A. Weis, Andersen Chang, Taliah Muhammad, Saumil Patel, Zhiwei Ding, Dat Tran, Jiakun Fu, Casey M. Schneider-Mizell, R. Clay Reid, Forrest Collman, Nuno Maçarico da Costa, Katrin Franke, Alexander S. Ecker, Jacob Reimer, Xaq Pitkow, Fabian H. Sinz, Andreas S. Tolias

**Affiliations:** Center for Neuroscience and Artificial Intelligence, Baylor College of Medicine, Houston, USA; Department of Neuroscience, Baylor College of Medicine, Houston, USA; Institute of Computer Science and Campus Institute Data Science, University Göttingen, Germany; Max Planck Institute for Dynamics and Self-Organization, Göttingen, Germany; Department of Electrical and Computer Engineering, Rice University, Houston, TX, USA; Institute for Bioinformatics and Medical Informatics, University of Tübingen, Germany; Allen Institute for Brain Science, Seattle, WA, USA; Department of Ophthalmology, Byers Eye Institute, Stanford University School of Medicine, Stanford, CA, US; Stanford Bio-X, Stanford University, Stanford, CA, US; Wu Tsai Neurosciences Institute, Stanford University, Stanford, CA, US; Department of Electrical Engineering, Stanford University, Stanford, CA, US

**Keywords:** Visual cortex, Foundation model, Generalization, Artificial Intelligence

## Abstract

The complexity of neural circuits makes it challenging to decipher the brain’s algorithms of intelligence. Recent breakthroughs in deep learning have produced models that accurately simulate brain activity, enhancing our understanding of the brain’s computational objectives and neural coding. However, these models struggle to generalize beyond their training distribution, limiting their utility. The emergence of foundation models, trained on vast datasets, has introduced a new AI paradigm with remarkable generalization capabilities. We collected large amounts of neural activity from visual cortices of multiple mice and trained a foundation model to accurately predict neuronal responses to arbitrary natural videos. This model generalized to new mice with minimal training and successfully predicted responses across various new stimulus domains, such as coherent motion and noise patterns. It could also be adapted to new tasks beyond neural prediction, accurately predicting anatomical cell types, dendritic features, and neuronal connectivity within the MICrONS functional connectomics dataset. Our work is a crucial step toward building foundation brain models. As neuroscience accumulates larger, multi-modal datasets, foundation models will uncover statistical regularities, enabling rapid adaptation to new tasks and accelerating research.

## Introduction

Recently, deep artificial neural networks (ANNs) have sparked a new era in building functional models of the brain that simulate brain activity based on sensory input, behavior, and internal states. For example, these models have set a new standard for functional models of the visual cortex (Yamins et al., 2014a; Cadieu et al., 2014; Antolík et al., 2016; Batty et al., 2017; McIntosh et al., 2016; Klindt et al., 2017; Kindel et al., 2017; Cadena et al., 2019; Burg et al., 2021; Lurz et al., 2020; Bashiri et al., 2021; Christensen and Zylberberg, 2020; Cowley and Pillow, 2020; Ecker et al., 2018; Sinz et al., 2018; Bakhtiari et al., 2021; Nayebi et al., 2021; Willeke et al., 2022). When these models are trained to optimize specific tasks such as object classification, next image prediction, or next word prediction, their hidden representations match those observed in the brain, providing a normative representational-level modeling approach to understanding the brain (Yamins et al., 2014b). With the collection of larger datasets in neuroscience, data-driven models have begun to outperform task-driven models (Pierzchlewicz et al., 2024). Whether they are data or task-driven, accurate functional models of the brain enable large-scale *in silico* experiments to be performed to systematically characterize neuronal representations and decipher principles that govern information processing in the brain. Because these models are differentiable, they allow for experiments like applying image synthesis methods, which are extremely difficult to perform in the brain without a model. For example, in vision, this approach can identify the most exciting stimulus for individual neurons (Bashivan et al., 2019; Walker et al., 2019; Ponce et al., 2019; Franke et al., 2022; Höfling et al., 2022), determine what individual neurons are selective for, characterize contextual modulation, explore how brain states affect tuning functions under natural stimulus conditions (Walker et al., 2019; Bashivan et al., 2019; Franke et al., 2022; Fu et al., 2023), and ascertain what they are invariant to (Ding et al., 2023b), aiming to move toward an interpretable understanding of the neural code. The resulting predictions can then be tested through *in vivo* closed-loop experiments, such as the *inception loops* paradigm (Walker et al., 2019; Franke et al., 2022; Bashivan et al., 2019). This *in silico*–*in vivo* approach addresses the inherent challenges of studying neuronal representations, including the high dimensionality of the input space, the nonlinear nature of information processing in the brain, and the limited availability of time for conducting *in vivo* experiments.

However, a challenge in neural network modeling is predicting new stimulus domains outside the original training distribution (Hendrycks and Dietterich, 2019). For instance, when models are trained to generate responses to natural movies, they perform well at predicting unseen natural movies but exhibit a substantial decrease in prediction performance on other domains such as synthetic or parametric stimuli (Sinz et al., 2018; Vystrčilová et al., 2024). However, to build upon the long history of using parametric stimuli for visual psychophysics and neurophysiology (Britten et al., 1992; Salzman et al., 1990; Marshel et al., 2019) and to increase their usefulness for *in silico* experiments, it is crucial to develop functional models that generalize well to novel stimulus domains, such that tuning functions can be characterized *in silico*, e.g., with parametric stimuli (Ustyuzhaninov et al., 2022). Recently, so called *foundation models* (Bommasani et al., 2021) in artificial intelligence, characterized by their ability to train on massive amounts of data and build robust representations of their modeling domain, have demonstrated remarkable generalization and capabilities in downstream tasks (Brown et al., 2020; Radford et al., 2021). For example, foundation models of language are trained on vast quantities of text encompassing much of human knowledge. Trained to predict the next sub-word in text, these foundation models capture robust language and knowledge representations that can be transferred to new tasks with relatively little data. These tasks include answering unstructured questions and even passing medical licensing exams (Kung et al., 2023).

Inspired by these breakthroughs, we sought to develop a foundation model of the mouse visual cortex trained on extensive quantities of data to predict neural activity from dynamic video and behavior as inputs. We collected the responses to ecological video stimuli from ∼135,000 neurons across multiple areas of the visual cortex from a total of 14 awake, behaving mice. Using a highly optimized deep neural network trained on a subset of these data from 8 mice, we learned a common, data-driven “foundation core” that effectively captured the shared latent representations of all recorded neurons which accurately predicted neuronal responses across many mice and visual cortical areas. New models utilizing the foundation core demonstrated the ability to be rapidly and accurately fitted to new mice with minimal amounts of data, surpassing the performance of individualized models that were trained end-to-end for each mouse individually. These models excelled not only in predicting neuronal responses to new natural movies (in-domain) but also generalized to accurately predict responses to various out-of-domain stimuli, including random moving dots, flashing dots, Gabor patches, coherent moving noise, and static natural images.

To test whether our foundation model could also be adapted to perform well in new tasks beyond neural activity prediction, we evaluated its performance in predicting morphologically defined cell types. We found that this model could accurately predict the anatomically defined cell type class of excitatory neurons from L2-5 in the multi-area MICrONS dataset (The MICrONs Consortium et al., 2023), which contains over 70,000 neurons within a ∼1mm^3^ cortical volume spanning multiple visual areas, featuring both nanoscale-level anatomical structure and responses to natural movies. Moreover, our model also predicted more specific morphological features of neurons, such as the dendritic bias of layer 4 exci-tatory neurons (Weis et al., 2022). Importantly, our model enabled a detailed analysis to characterize the relationship between synaptic level connectivity and the functional properties of neurons (Ding et al., 2023a), including tuning invariance (Ding et al., 2023b) and contextual modulation (Fu et al., 2023).

As extensive multi-modal neuroscience datasets accumulate, we anticipate the emergence of powerful foundational brain models, akin to large language models. These models will unveil statistical patterns spanning sensory, behavioral, and activity modalities. By integrating with molecular and anatomical foundation models, they will not only advance fundamental neuroscience but also pave the way for innovative disease treatments.

## Results

### State-of-the-art dynamic functional model of the mouse visual cortex

To model the dynamic neuronal responses of the mouse visual cortex, we developed an ANN that was comprised of four modules: perspective, modulation, core, and readout (Fig. 1). The modular design enabled the ANN to accommodate diverse tasks and inputs. For instance, eye movements and different positioning of a mouse’s head relative to the monitor can result in different perspectives of the same stimulus, despite best efforts to limit experimental variability. To account for this, the perspective module of our ANN uses ray tracing and eye tracking data to infer the perspective of the mouse from the presented stimulus on the monitor (Extended Data Fig. 1). To account for behavioral factors that modulate the activity of the visual cortex (Reimer et al., 2014), the modulation module transforms behavioral inputs (locomotion, pupil dilation) to produce dynamic representations of the mouse’s behavioral and attentive state (Extended Data Fig. 2). The perspective and modulation modules provide visual and behavioral inputs, respectively, to the core module of the ANN. Composed of feedforward (3D convolution layers) and recurrent (long-short term memory) components, the core contains the majority of the ANN’s modeling capacity and produces nonlinear representations of vision that are modulated by behavior. These representations are mapped onto the activity of individual neurons by the readout module, which performs a linear combination of the features generated by the core at one specific location, the neuron’s receptive field. All four modules of the ANN (perspective, modulation, core, and readout) were trained end-to-end to predict time series of neuronal responses to natural movies (for details of model architecture and training, see Methods).

**Fig. 1.**
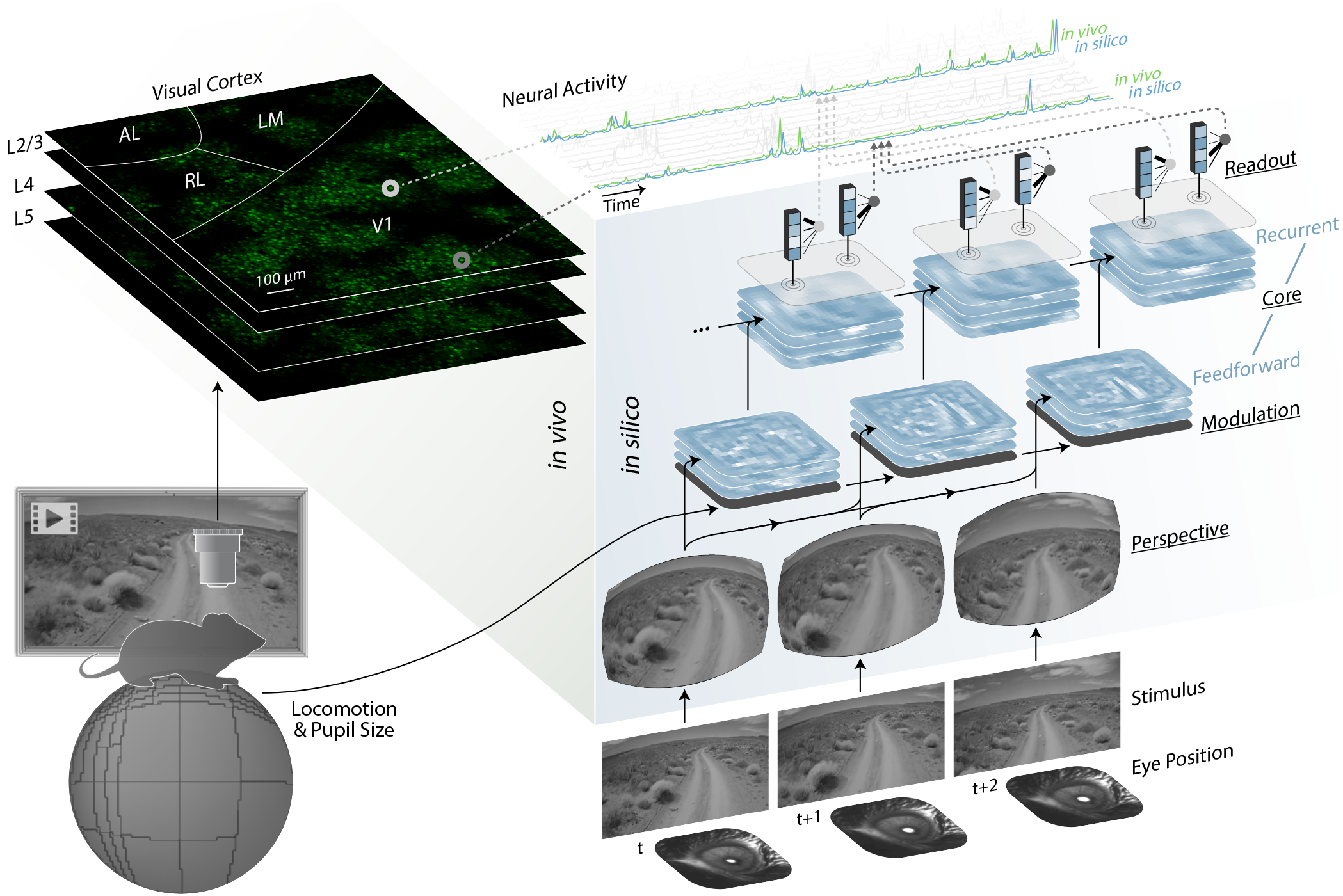
ANN model of the visual cortex. The left panel (green) depicts an *in vivo* recording session of excitatory neurons from several areas (V1, LM, RL, AL) and layers (L2/3, L4, L5) of the mouse visual cortex. The right panel (blue) shows the architecture of the ANN model and the flow of information from inputs (visual stimulus, eye position, locomotion, and pupil size) to outputs (neural activity). Underlined labels denote the four main modules of the ANN: perspective, modulation, core, and readout. For the modulation and core, the stacked planes represent feature maps. For the readout, the blue boxes represent the core’s output features at the readout position of the neuron, and the fanning black lines represent readout feature weights. The top of the schematic displays the neural activity for a sampled set of neurons. For two example neurons, *in vivo* and *in silico* responses are shown (green and blue, respectively).

First, we evaluated the predictive accuracy of our ANN model architecture when trained on on individual recording sessions lasting ∼1 hour. Predictive accuracy was measured by the correlation between the recorded and the predicted responses to a novel set of stimuli that were not included in model training. To account for *in vivo* noise, the correlation was normalized by an estimated upper bound on the performance that could be achieved by a perfect model (Schoppe et al., 2016). Using this normalized correlation coefficient (*CC*_norm_) as the metric of predictive accuracy, we compared our model to the previous best-performing dynamic model of the mouse visual cortex (Sinz et al., 2018). Trained and tested on the same data from that study (dynamic V1 responses to natural movies), our model had a 25–46% increase in predictive accuracy on held-out test data across the three recording sessions used in Sinz et al. 2018 (Fig. 2a). This level of increase in performance is substantial for predictive models of the visual cortex. We also evaluated the predictive accuracy of our model on newly collected data that contained multiple visual areas (Fig. 2b). Interestingly, we found that the performance of our model for higher visual areas (LM, RL, AL) was similar to V1 (Fig. 2c), despite the increased complexity of neuronal tuning to more complex features exhibited by higher visual areas (Siegle et al., 2021; Goltstein et al., 2021). Next, we performed lesion studies to determine the effect that individual components of the model had on predictive accuracy (Extended Data Fig. 3). Removing either of the two behavioral modules resulted in a modest but significant reduction in reduced predictive accuracy: 2.3% reduction for the perspective module (Extended Data Fig. 3a–e) and 2.8% for modulation module (Extended Data Fig. 3f–j). For the core component, we found that using 3D convolutions in the feedforward component significantly improved performance compared to 2D convolutions, although the difference was small at 0.88% (Extended Data Fig. 3k–o). We also evaluated the objective function used for training and found that the Poisson negative log likelihood loss significantly outperformed mean squared error loss, with a performance difference of 9.6% (Extended Data Fig. 3p–t). In summary, our new ANN model sets new standards for predicting dynamic neuronal responses of the visual cortex, with individual components contributing modest but significant improvements. Importantly, the main driver of increased performance is the much larger dataset used for training, aligning with scaling laws observed in AI research - a property exhibited by ANNs in general where performance improves with increased data (Fig. 2b).

**Fig. 2.**
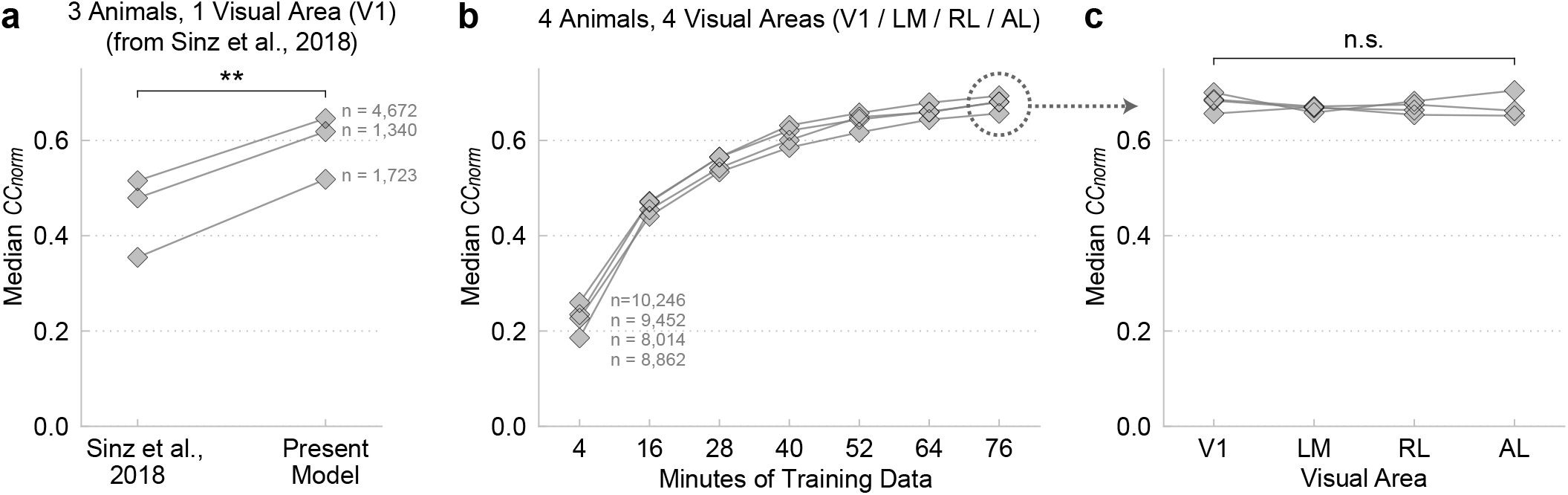
Predictive accuracy of models trained on individual recording sessions. **a**, Predictive accuracy (median *CC*_norm_ across neurons, see Methods for details) of our model vs. the previous state-of-the-art dynamic model of the mouse visual cortex by Sinz et al. (2018). We trained and tested our model on the same set of data from Sinz et al. (2018): V1 neuronal responses to natural movies from 3 mice. n = number of neurons per mouse. ** = paired two-way *t* -test, t=14.53, p < 0.01, df=2. **b**, Predictive accuracy of our models by the amount of data used for training for 4 new recording sessions and mice. For each recording session, training data was partitioned in to 7 fractions ranging from 4 to 76 minutes. Separate models (diamonds) were trained on the differing fractions of training data, but tested on the same held-out testing data. Models of the same mice are connected by lines. **c**, Predictive accuracy by visual area, from models that were trained on the full data. We did not find a statistically significant relationship between predictive accuracy and visual areas (linear mixed effects model (Lindstrom and Bates, 1988), n.s. = Wald test, p = 0.45, df=3).

### Foundation models generalize to new subjects and stimulus domains

The remarkable performance of foundation models in other domains—e.g., natural language (Brown et al., 2020) and image generation (Radford et al., 2021)— originates from their vast quantities of training data. How-ever, collecting large amounts of neuronal data from individual neurons and animals presents challenges. Individual recording sessions are limited in duration by experimental factors such as attentiveness and recording device stability. To overcome this limitation, we combined data from multiple recording sessions, totaling over 900 minutes of natural movie responses from 8 mice, 6 visual areas (V1, LM, AL, RL, AM, PM), and ∼66,000 neurons. This data was used to train a single, shared ANN core (Fig. 3a) with the goal of capturing common representations of vision that underlie the dynamic neuronal response of the visual cortex for a representative set of neurons and a group of mice. This representation could then be used to fit models of new mice to improve their performance with limited data. Here we refer to the representative group of 8 mice as the “foundation cohort”, the trained ANN component as the “foundation core”, and ANNs derived from the foundation core as “foundation models”.

**Fig. 3.**
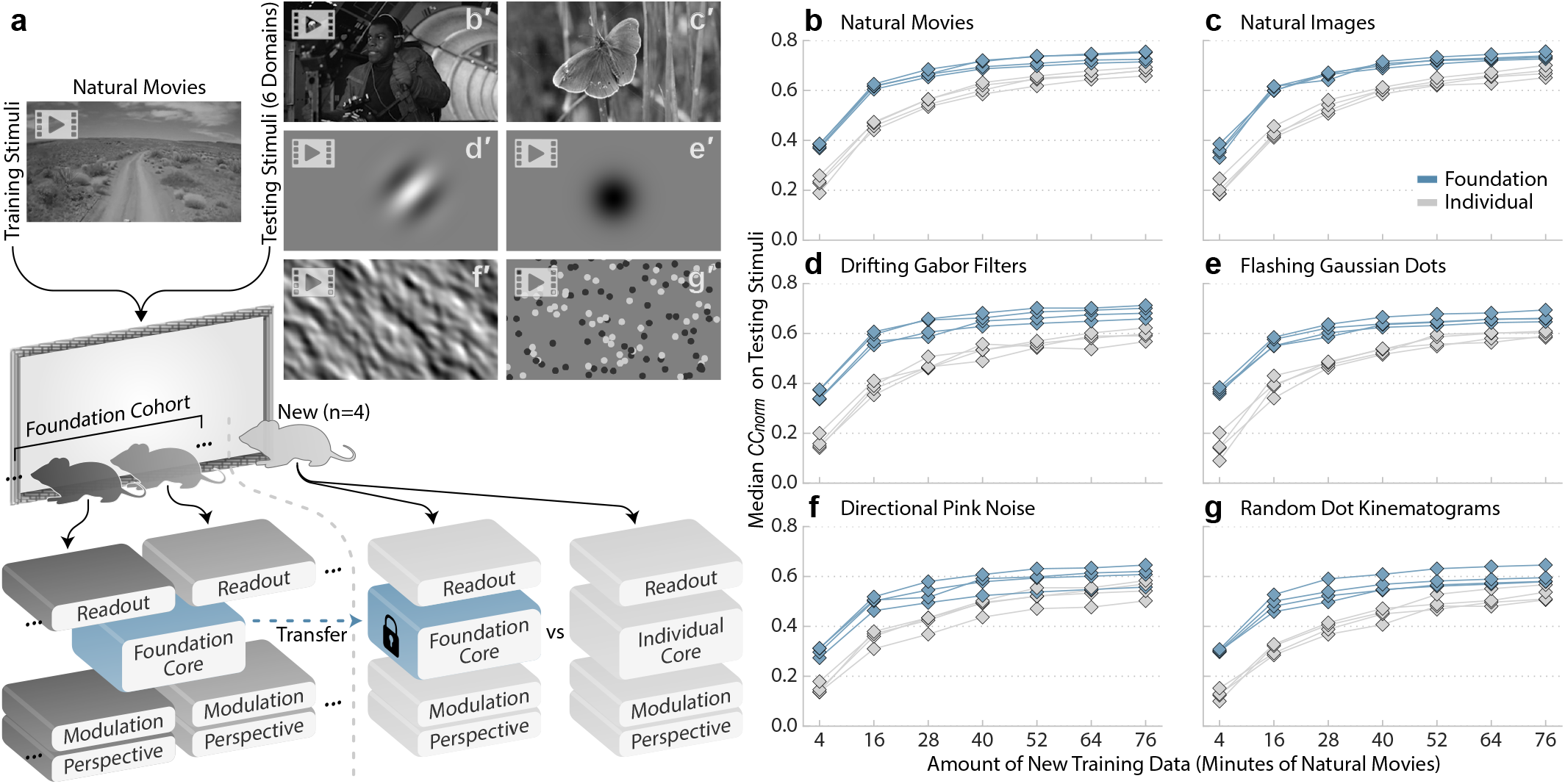
Predictive accuracy of foundation models. **a**, Schematic of the training and testing paradigm. Natural movie data were used to train: 1) a combined model of the foundation cohort of mice with a single foundation core, and 2) foundation models vs. individual models of new mice. The models of the new mice were tested with stimuli from 6 different domains (**b’**–**g’**). **b**–**g**, Corresponding plots show the predictive accuracy (median *CC*_norm_ across neurons) as a function of the amount of training data for foundation models (blue) vs. individual models (gray) of the new mice. 4 mice × 7 partitions of training data ×2 types of models = 56 models (diamonds). Models of the same mouse and type (foundation / individual) are connected by lines. Number of neurons per mouse = 8,862 | 8,014 | 9,452 | 10,246.

To evaluate the representation of the visual cortex captured by the foundation core, we froze its parameters and transferred it to ANNs with new perspective, modulation, and readout components fitted to new mice (Fig. 3a). Each new mouse was shown an assortment of stimuli, designated for either model training or testing. The training stimuli consisted of natural movies, and we used different portions of this, spanning from 4 to 76 minutes, to fit ANN components to the new mice. This approach aimed to examine the relationship between the models’ performance and the amount of training data for each new mouse. The testing stimuli included natural movies that were not part of the training set (Fig. 3b’), and new stimulus domains like static natural images (Fig. 3c’), and 4 types of parametric stimuli (Fig. 3d’–g’), consisting of drifting Gabor filters, flashing Gaussian dots, directional pink noise, and random dot kinematograms. To test the role of the foundation core in prediction performance, we trained a set of control models that differed from the foundation models only by the core component. For these controls or “individual models”, all four components—core, perspective, modulation, and readout—were trained end-to-end using training data from a single recording session. For the foundation models, training data from the new mice were only used to fit the perspective, modulation, and readout components, and the core was trained on the foundation cohort as described above and was frozen (Fig. 3a).

When tested on natural movies, foundation models outperformed individual models and required less training data from the new mice to achieve high levels of predictive accuracy (Fig. 3b). For instance, individual models required more than an hour of training data to surpass a median *CC*_*norm*_ of 0.65 for all mice, whereas foundation models required less than half an hour (Fig. 3b). This performance gain was observed across all tested stimulus domains, including those that were in new stimulus domains (Fig. 3c’–g’), i.e., outof-distribution (OOD) from the training domain of natural movies (Fig. 3b’). Importantly, no stimuli from the OOD domains were used to train any component of the models, including the foundation core. Nevertheless, foundation models were more accurate at predicting responses to new stimulus domains while requiring substantially less training data from the new mice (Fig. 3c–g). For example, when predicting drifting Gabor filters, the foundation models were able to achieve a performance of median *CC*_*norm*_ *>* 0.55 using only 16 minutes of natural movie training data. In contrast, the individual models required more than an hour of training data to reach the same performance level (Fig. 3d). This highlights the significant difference in the data efficiency of these models, i.e., the amount of training data (sample complexity) required from new subjects to accurately fit their neuronal responses. Thus, training a foundation dynamic core on natural movie data pooled from multiple cortical layers, areas, and animals produces a robust and transferable representation of the visual cortex that generalizes to new animals and improves model performance for not only natural movies but also novel stimulus domains.

When combining functional studies of the brain with other modalities like anatomy, there is typically a limited amount of time available for *in vivo* recordings before destructive histological analysis is performed. While traditionally this would limit the number of functional studies that can be performed *in vivo*, predictive models allow essentially unlimited scans to be performed *in silico*, even after tissue has been destroyed. To enable this for the MICrONS project, responses to natural movies were collected for the purpose of model training. Due to the challenge of completing all 14 scans in the same animal in as short a period as possible, the amount of training data collected from each experiment (mean 42 minutes, range 33–53 minutes, depending on optical quality and animal behavioral profile) was less than the other recording sessions in this paper. With the available amount of data, individual models—with all components trained on a single experiment—achieved a median *CC*_norm_ of 0.48–0.65, when tested on a held-out set of natural movies. By applying our foundation modeling paradigm—transferring the foundation core and fitting only the perspective, modulation, and readout components on a single experiment—the median *CC*_norm_ increased to 0.58–0.76 (Extended Data Fig. 4). This highlights the advantage of the foundation modeling approach when there is a limited amount of data available for training.

### Foundation models enable classical studies of parametric tuning

By leveraging the foundation core and transfer learning, we were able to create accurate foundation models for individual mice (Fig. 3). These models enable essentially unlimited *in silico* experiments for studying representations, testing theories, and generating novel hypotheses that can be verified *in vivo*. Here we assessed the precision with which classical tuning properties of the visual cortex could be replicated at the individual neuronal level in our foundation model. We presented mice—not part of the foundation cohort—with natural movie stimuli in order to train their ANN counterparts (Fig. 4a). Additionally, we presented parametric stimuli (Fig. 4b’–c’) to measure the orientation, direction and spatial tuning of the recorded neurons. Subsequently, we presented the same parametric stimuli to the corresponding *in silico* neurons and measured their properties for comparison (Fig. 4b–c). This was done for 3 mice and ∼30,000 neurons from 4 visual areas (V1, LM, AL, RL). To measure orientation and direction tuning, we presented directional pink noise (Fig. 4b’), which encoded coherent motion of different directions (0–360°) and orientations (0– 180°). First, we computed the strength of orientation and direction tuning via selectivity indices for orientation (OSI) and direction (DSI). There was a high correspondence between *in vivo* and *in silico* estimates for both OSI (Fig. 4d) and DSI (Fig. 4f), which validated the foundation model’s estimates of tuning strength for orientation and direction. Next, we estimated the preferred angles of orientation and direction of neurons by fitting a directional parametric model (mixture of von Mises distributions) to the responses. For strongly tuned neurons, the *in vivo* and *in silico* estimates of preferred angles of orientation and direction were closely matched (Fig. 4e,g). For example, for strongly orientation-tuned neurons with an *in silico* OSI > 0.5 (11% of neurons), the median difference between the *in vivo* and *in silico* estimates of preferred orientation was 4°, and with a lower OSI threshold of > 0.3 (43% of neurons), the median difference was 7°(Fig. 4e).

**Fig. 4.**
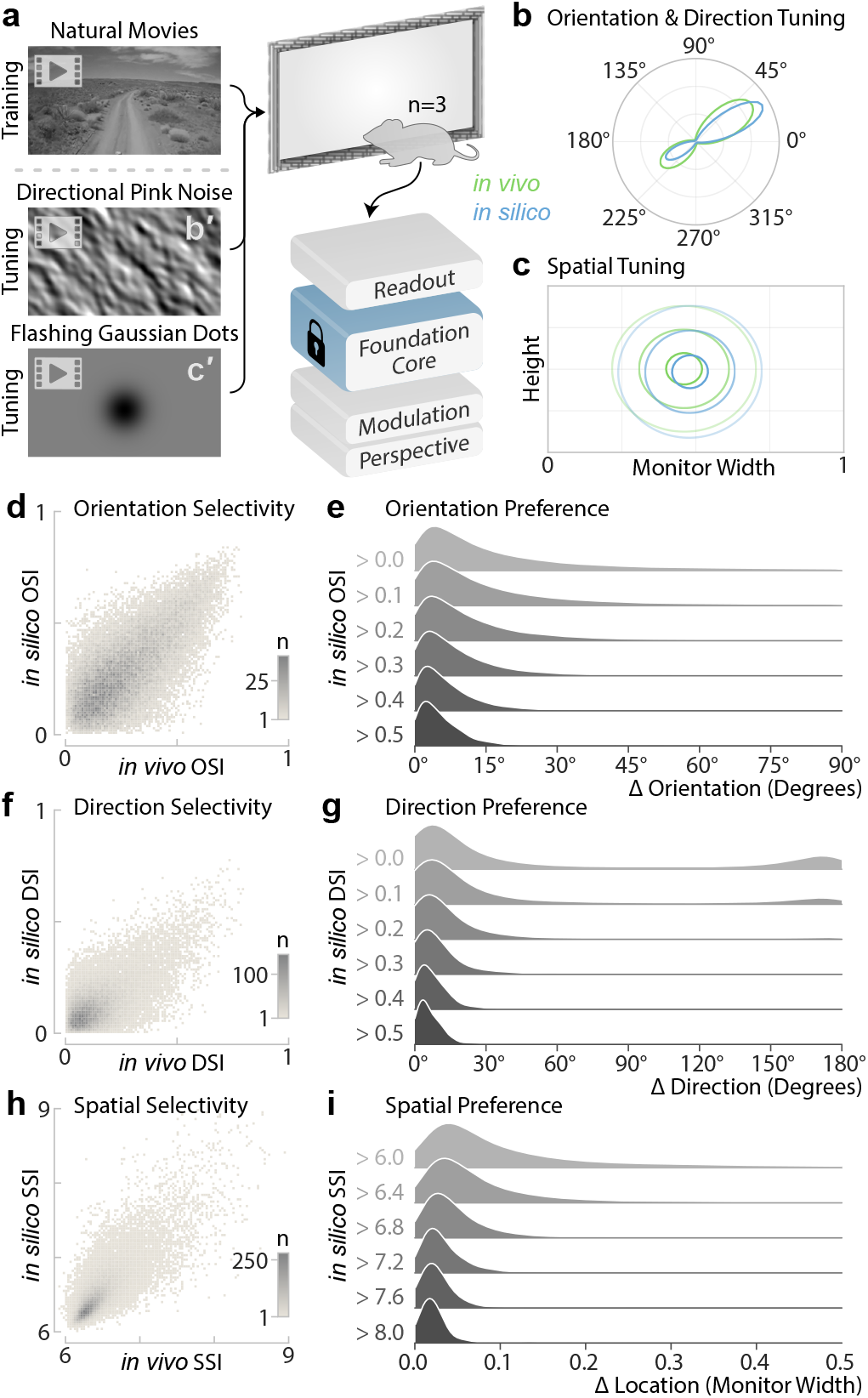
Parametric tuning from the foundation model. **a**, Schematic of the experimental paradigm: foundation models of new mice (n=3) were trained with natural movies, and estimates of parametric tuning were computed from *in vivo* and *in silico* responses to synthetic stimuli (**b’**, directional pink noise; **c’**, flashing Gaussian dots). **b**,**c**, *In vivo* and *in silico* estimates of an example neuron’s parametric tuning to orientation/direction (**b**) and spatial location (**c**). **d**,**f**,**h**, Binned scatter plots of *in vivo* and *in silico* estimates of selectivity indices (SI) for orientation (**d**, OSI), direction (**f**, DSI), and spatial (**h**, SSI). The color indicates the number of neurons (n) in each bin. **e**,**g**,**i**, Density histograms of differences between *in vivo* and *in silico* estimates of preferred orientation (**e**), direction (**g**), and spatial location (**i**). In each panel, histograms containing increasingly selective groups of neurons, thresholded by *in silico* OSI (**e**) / DSI (**g**) / SSI (**i**), are stacked from top to bottom. The density histograms were produced via kernel density estimation using Scott’s bandwidth.

To measure spatial tuning, we presented flashing Gaussian dots (Fig. 4c’) to the neurons described above. We computed a spike-triggered average (STA) of the stimulus, which was used to estimate: 1) the strength of spatial tuning for Gaussian dots (non-uniformity of the STA) via the spatial selectivity index (SSI); and 2) the preferred location (peak of the STA) via least squares fitting of the STA to a spatial parametric model (2D Gaussian distribution). Although using the Gaussian dot stimulus did not elicit strong SSI for the majority of neurons, for those neurons that were strongly tuned *in silico*, we observed a close match between *in vivo* and *in silico* estimates of spatial tuning strength, measured by SSI (Fig. 4h). For instance, for strongly tuned neurons with *in silico* SSI > 8, the median distance between the *in vivo* and *in silico* estimates of the preferred location was 0.02 of the monitor width (Fig. 4i), approximately 2° in visual space.

Together, these results demonstrate the accuracy of estimating tuning parameters for classical functional properties from our foundation model with no prior training on parametric stimuli. Therefore, rather than presenting parametric stimuli *in vivo*, parametric tuning can be performed *in silico* with an accurate and validated foundation model, freeing up valuable *in vivo* experimental time for other purposes.

### Foundation model predicts structural properties of neurons in the MICrONS dataset

The function of the neocortex mechanistically emerges from its circuit structure. The MICrONS project, a landmark dataset in neuroscience, provides unprecedented scale and resolution, combining millimeter-scale functional data with structural data that spans a similar volume but at nanometer resolution, across multiple visual cortical areas of a single mouse. In the MICrONS mouse, the responses of over 70,000 excitatory neurons to natural movies were measured across 14 sequential scans, encompassing a 1 cubic millimeter volume spanning V1, LM, AL, and RL visual areas. This volume was subsequently subjected to serial electron microscopy (EM) and dense morphological reconstruction (Fig. 5b), resulting in detailed structures of approximately 60,000 excitatory neurons and 500 million synapses, representing the largest integrated study of neocortical structure and function to date (The MICrONs Consortium et al., 2023).

**Fig. 5.**
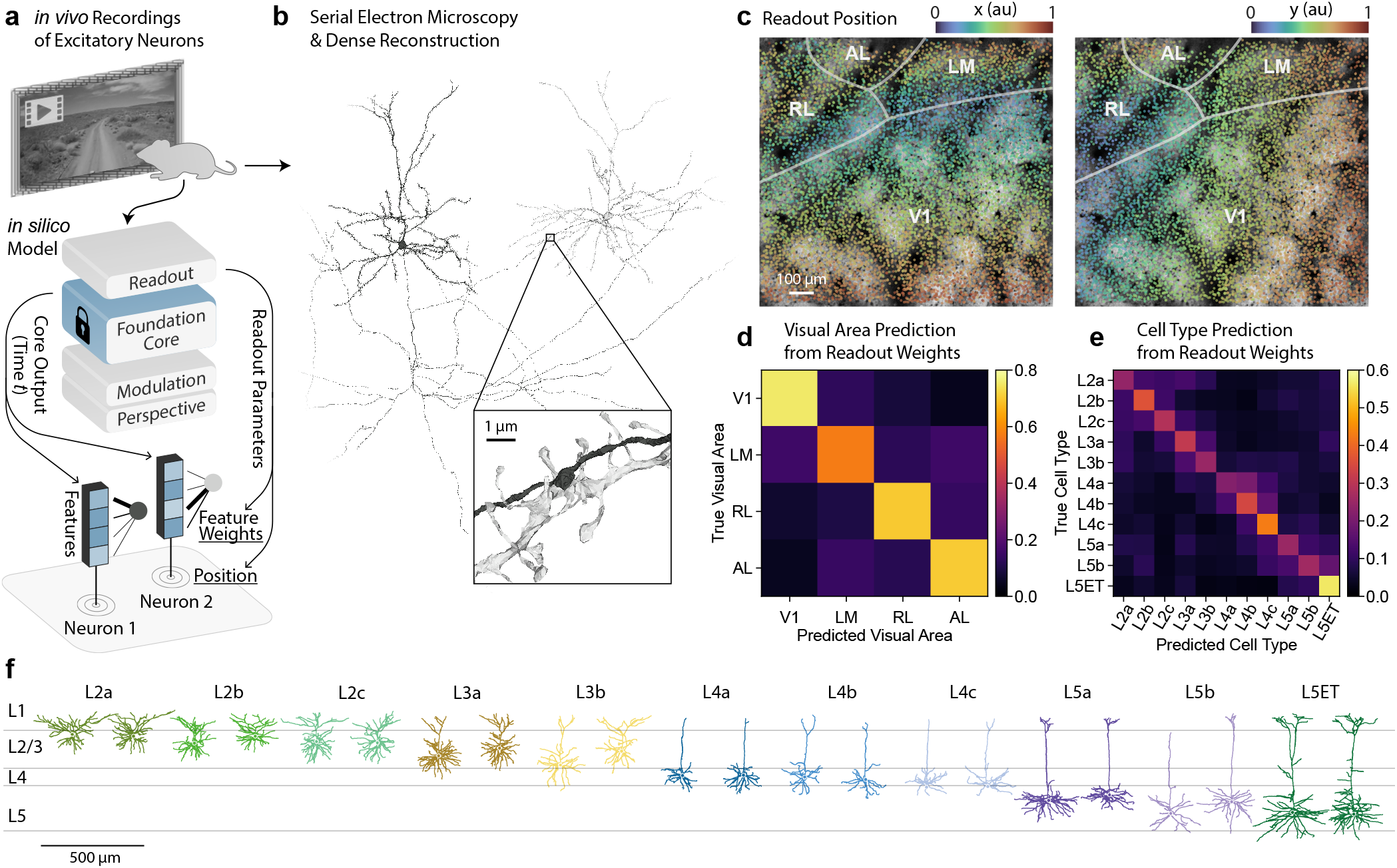
The foundation model of the MICrONS volume relates neuronal function to structure and anatomy. a, Schematic of a foundation model of the MICrONS mouse, trained on excitatory neuronal responses to natural movies. At the bottom, the readout at a single time point is depicted, showing the readout positions and feature weights for two example neurons. b, Meshes of two example neurons, reconstructed from serial electron microscopy. The zoom-in cutout shows a synapse between these two neurons, with the pre-synaptic axon in black and post-synaptic dendrite in silver. c, Colored scatter plots of readout positions of all neurons from a recording session of the MICrONS mouse, overlayed on top-down a view of the recording window with annotated visual areas (V1, LM, RL, AL) and boundaries. The left and right plots are colored by the x and y coordinates of the readout positions, respectively. d, Confusion matrix of MICrONS visual areas predicted from readout feature weights, normalized per row. The diagonal represents the recall for each visual area. e, Confusion matrix of MICrONS excitatory neuron cell types predicted from readout feature weights, normalized per row. The excitatory neuron cell types are from Schneider-Mizell et al. 2023. The diagonal represents the recall for each cell type. f, Morphologies of different types excitatory neurons. Two example neurons are shown for each excitatory neuron cell type.

We employed the foundation modeling paradigm to the MI-CrONS dataset to model the function of excitatory neurons within the 1 cubic millimeter volume. The model’s read-out module maps the foundation core’s output onto individual neuronal responses. Each neuron’s readout parameters consist of two components: readout position and readout feature weights (Fig. 5a). We trained readout parameters for all excitatory neurons recorded in the MICrONS volume, and we investigated whether these parameters would be useful for studying the structure-function relationship of the brain.

We first examined the readout position, which consists of two parameters per neuron: azimuthal (x) and altitudinal (y) locations, specifying the center of the receptive field learned by the model for each neuron. Analysis of the readout positions revealed that they accurately captured the retinotopic organization of the visual cortex (Fig. 5c). In V1, readout x positions aligned with the medial-lateral axis, and y positions aligned with the rostral-caudal axis. At the border of V1 and LM/RL, there was an inversion of the axis for the x readout position, demarcating the transition zone between these areas. This organization of readout positions according to anatomical locations aligns well with prior studies of retinotopic organization in the mouse visual cortex (Garrett et al., 2014; Zhuang et al., 2017).

Next, we investigated how the readout weights, a 512-dimensional vector per neuron, could be used to predict anatomical properties such as the visual area and morphologically defined cell types. These readout weights serve as a functional barcode, encoding the neuron’s tuning to visual features produced by the core module at its readout position. We found that these functional barcodes captured differences between visual areas (V1, LM, AL, RL). Using logistic regression, the readout weights could predict visual areas with a balanced accuracy of 68%, exceeding the chance level of 25% (Fig. 5d). We further explored the possibility of predicting 11 morphologically defined excitatory cell types from L2 to L5 (Fig. 5f) identified by Schneider-Mizell et al. 2023. Again, employing logistic regression, we achieved a balanced accuracy of 32% for cell type prediction, outperforming the chance baseline of 9% (Fig. 5e). Because these cell types are fairly well separated across cortical depth (Fig. 5f), it is possible that the classifier has learned to predict depth directly from the depth-varying signal-to-noise ratio of twophoton (2p) imaging. To control for this potential confound, we trained a classifier to predict cell types from 2p depth (re-duced model) and compared to a second classifier provided with both 2p depth and readout feature weights (full model). We found that the full model significantly outperformed the reduced model in predicting cell types (likelihood ratio test, p < 10^−9^), indicating that the readout feature weights contribute to classifier performance. Collectively, these results demonstrate that our foundation model captures both functional and structural properties of neurons, making it a valuable tool for analyzing structure-function relationships within the MICrONS volume and studying mechanisms of computation within the visual cortex.

Our foundation model was employed in a companion paper (Ding et al., 2023a) to investigate the wiring rules of mouse visual cortex. The model’s ability to factorize each neuron’s visual tuning into spatial (‘where’) and feature (‘what’) components—encoded by readout position and readout weights, respectively—enabled a novel analysis of function-connectivity relationships. Their study revealed that the feature component, rather than the spatial component, was predictive of synaptic connectivity between neurons. This approach extends beyond previous studies that relied on signal correlation, as our model separates feature selectiv-ity and receptive field location from underlying visual scene statistics. Consequently, it provides a more nuanced understanding of how specific neuronal characteristics influence connectivity patterns in the visual cortex. In another companion study, it was used to investigate a unique morphological variation of neurons in layer 4 of the MICrONS dataset (Weis et al., 2022). This study discovered a subset of excitatory neurons at the base of layer 4 with restricted dendritic reach, quantified by a basal dendritic bias metric. Remarkably, our functional barcodes accurately predicted this basal dendritic bias (Weis et al., 2022). This capability bridges the traditional divide between functional properties and morphological traits in neuroscience, enabling exploration of structurefunction relationships in cortical circuits. Thus, our foundation model and the functional digital twin of the MICrONS mouse represent a significant advance in linking neuronal anatomy and physiology.

## Discussion

We introduce a major step towards a foundation model for the mouse visual cortex that achieves state-of-the-art performance at predicting dynamic neuronal responses across multiple visual areas, marking significant progress towards an accurate functional digital twin of the mouse visual system.

Beyond excelling in the natural movie domain on which it was trained, it accurately predicted responses to new stimulus domains, including coherent random moving dots, dynamic Gabor patches, flashing dots, directional pink noise, and natural static images. The model’s generalization performance on new stimulus domains highlights its ability to capture nonlinear transformations from image space to neuronal activity in the mouse visual cortex. The foundation core enabled accurate models of new mice to be fitted with limited training data, outperforming models with cores that were individually trained for each mouse underscoring the inter-individual similarity between individual mice (Lurz et al., 2021).

Importantly, we also demonstrate the utility of our model for making predictions beyond neural activity—for example, in tasks related to anatomy and connectivity—which greatly enhances its utility as a foundation model of the brain (Bommasani et al., 2021). Specifically, using the foundation core of our model, we built a functional digital twin of the MICrONS mouse functional connectomics dataset. No anatomical information from EM data was used to build this model. The functional digital twin enabled us to extract a functional barcode—a vector embedding that describes the input-output function of each neuron. Using these functional barcodes, we show that our model could predict the cell type identity of these neurons, which were defined by morphological and EM characteristics in a companion paper analyzing the MICrONS anatomical data (Schneider-Mizell et al., 2023). Moreover, in an additional companion paper characterizing the morphological landscape of the MICrONS dataset, our model was used to predict specific features of the dendritic morphology of layer 4 pyramidal neurons (Weis et al., 2022).

In a companion paper, our digital twin of the MICrONS mouse was used to analyze the relationship between neuronal function and connectivity (Ding et al., 2023a). This was made possible by the architecture of the foundational model, which allowed the tuning properties of each simulated neuron to be represented as a functional barcode (indicating what the neuron responds to) and a spatial component (indicating the position of the neuron’s receptive field). This factorization of the neuronal tuning functions was then leveraged by Ding et al. 2023a to examine the relationship between the functional properties of neurons and their synaptic connectivity at an unprecedented level of detail. The key finding from this study was that the feature component of neuronal tuning, but not the spatial component, predicted which neurons were connected at the fine synaptic scale.

Another important utility of our foundation model of the mouse visual cortex, and the digital twin built from the MICrONS dataset, is that the digital twin effectively “immortalizes” the functional properties of the recorded neurons in the study. If the foundation core can demonstrate generalization to novel stimulus domains of interest, the digitally twinned neurons can be characterized using these new stimuli that were not presented at the time of the *in vivo* data collection. Here, we demonstrate that models trained on natural movies indeed generalize to novel stimulus domains such as coherent random dots, noise patterns and static natural images. Naturally, conducting new animal validation experiments, similar to those presented in this study, will be necessary to confirm that the foundation core indeed generalizes to other animals and new stimulus domains of interest. With proper validation, a foundation model enhances the utility of the MICrONS dataset for broader impact since one can study specific functional properties of interest and determine how they are related to the microcircuit and anatomy of the neocortex.

To this end, Fu et al. 2023 performed extensive validation of our MICrONS model and utilized image synthesis to study the circuit wiring that could explain center-surround receptive field properties of neurons. They found that neurons neurons with similar feature selectivity, measured with our functional barcodes, were equally likely to form excitatory synapses, regardless of whether their receptive fields significantly overlapped or had minimal overlap. Deciphering this relationship between neuronal function and connectivity was made possible with our foundation model.

In summary, the results presented here and in the companion papers (Ding et al., 2023a; Weis et al., 2022; Fu et al., 2023) utilizing our model demonstrate the power of the foundation modeling approach for neuroscience research. In addition to achieving excellent neural predictions for not only new mice but also new stimulus domains, our model enabled the discovery of more nuanced synaptic connectivity rules and the prediction of morphological cell types, dendritic features, and related center-surround contextual modulation to local circuit wiring—all from the functional barcodes of our foundational model. This ability to uncover subtle patterns in neural organization showcases the model’s potential for driving new insights in neuroscience. In large projects like MICrONS, where dataset longevity is highly desirable, the strong generalization capabilities of foundation models and their ability to perform tasks beyond their original training offer clear benefits. This enables researchers to explore questions that were not necessarily designed into the original experiment, extending the utility of the dataset beyond its initial scope and facilitating novel discoveries in neural circuit organization.

Our work was inspired by recent breakthroughs in artificial intelligence, where foundation models (Bommasani et al., 2021), trained on massive data volumes, have demonstrated remarkable generalization in many downstream tasks. For example, models trained on next sub-word prediction (Brown et al., 2020) can be transferred to downstream tasks—e.g., conversing naturally with humans, or passing professional licensing exams (Kung et al., 2023)—with relatively small amounts of new data. Applied to neuroscience, the foundation modeling paradigm overcomes a major limitation of previous common approaches where models are individually trained using data from a single experiment. The limited amount of data hinders the accuracy of models as they try to learn from scratch the complex non-linearities of the brain, even though there is a great deal of similarity in how visual neurons respond. By contrast, foundation models combine data from multiple experiments, including data from many brain areas and subjects under high entropy natural conditions, giving them access to a much larger and richer set of data; only the specific idiosyncrasies of each individual mouse and its neurons must be learned separately. In other words, the similarities between neurons and subjects can be leveraged to identify common features of the brain, producing a more unified and accurate model of the brain that is informed by multiple subjects rather than one.

In neuroscience, previous work (Lurz et al., 2021) has shown that static models of the visual cortex benefit from pretraining on large amounts of data pooled from multiple subjects. In our current study, we not only demonstrate that data pooling and transfer learning can extend to a dynamic model, but we also crucially show that our model predicts neuronal responses to new stimulus domains (e.g., random dots and noise patterns) and new tasks such as anatomical features and connectivity, which are significant extensions over previous work. These findings and capabilities of large data-driven models of the brain underscore the potential of brain foundation models to study complex biological systems such as the brain.

Our present foundation model merely scratches the surface, as it only models parts of the mouse visual system under passive viewing conditions. By expanding this approach to encompass complex, natural behaviors in freely-moving subjects, incorporating additional brain regions, diverse cell types, and creating foundation models for other species could be a paradigm shift in neuroscience. Foundation models can be employed to study vision, cognition and motor control during intricate, unconstrained natural behaviors in which identical conditions rarely occur twice. For instance, we can conduct comprehensive *in silico* experiments to explore relationships between the high dimensional neuronal activity and behavioral spaces to generate hypotheses and to design simpler experiments to run *in vivo*, such as inception loops (Walker et al., 2019; Franke et al., 2022).

Moreover, by considerably reducing the neuron-hours required to model new individuals and behaviors, foundation models facilitate more efficient and cost-effective neuroscience experiments. For example, we can establish highthroughput research platforms that, with minimal new data, generate predictions of individual subjects’ neuronal activity and behavior. When causal manipulations are incorporated in the foundation model, such as pharmacological interventions, we could then swiftly screen drugs tailored for a desired phenotypic neuronal or behavioral outcome.

Ultimately, the development of multimodal foundation neuroscience models offers a powerful new approach to deciphering the algorithms underpinning natural intelligence. As we accumulate more diverse multimodal data—encompassing sensory inputs, behaviors, and neural activity across various scales, modalities and species—we will build powerful foundation models. This approach holds the promise of cracking the neural code of natural intelligence, providing unprecedented insights into the fundamental principles of cognition.

## ACKNOWLEDGEMENTS

The authors thank David Markowitz, the IARPA MICrONS Program Manager, for his support during all three phases of the MICrONS program. We thank IARPA program managers Jacob Vogelstein and David Markowitz for co-developing the MICrONS program. We thank Jennifer Wang, IARPA SETA for her assistance.

The work was supported by the Intelligence Advanced Research Projects Activity (IARPA) via Department of Interior/ Interior Business Center (DoI/IBC) contract numbers D16PC00003, D16PC00004, and D16PC0005. The U.S. Government is authorized to reproduce and distribute reprints for Governmental purposes notwithstanding any copyright annotation thereon. XP and AST acknowledge support from NSF NeuroNex grant 1707400. AST also acknowledges support from the National Institute of Mental Health and National Institute of Neurological Disorders And Stroke under Award Number U19MH114830 and National Eye Institute award numbers R01 EY026927 and Core Grant for Vision Research T32-EY-002520-37. Disclaimer: The views and conclusions contained herein are those of the authors and should not be interpreted as necessarily representing the official policies or endorsements, either expressed or implied, of IARPA, DoI/IBC, or the U.S. Government.

ASE received funding from the European Research Council (ERC) under the European Union’s Horizon Europe research and innovation programme (Grant agreement No. 101041669) as well as the Deutsche Forschungsgemeinschaft (DFG, German Research Foundation), project ID 432680300 (SFB 1456, project B05)

We also would like to thank Matthias Bethge, Mackenzie Mathis, Blake Richards, Anthony Zador and Joel Zylberberg for many stimulating discussions regarding building foundation models for the brain.

## AUTHOR CONTRIBUTIONS

We adopted the following contribution categories from CRediT (Contributor Roles Taxonomy). Authors within each category are sorted in the same order as in the author list.

**Conceptualization**: EYW, AST

**Methodology**: EYW, FHS, AST

**Software**: EYW, FHS

**Validation**: EYW, PGF, ZhuoD, ZhiD, DT, JF

**Formal analysis**: EYW, StP, AC

**Investigation**: EYW, PGF, KP, TM

**Resources**: PGF, SaP, StP

**Data Curation**: PGF, StP

**Writing - Original Draft**: EYW, PGF, StP, FHS, AST

**Writing - Review & Editing**: EYW, PGF, StP, KF, ASE, JR, XP, FHS, AST

**Visualization**: EYW, StP

**Supervision, Project administration, and Funding acquisition**: AST

## Methods

### Neurophysiological experiments

MICrONS data in Fig. 5 was collected as described in The MICrONs Consortium et al. 2023, and data in Fig. 2a was collected as described in Sinz et al. 2018. Data collection for all other figures is described below.

All procedures were approved by the Institutional Animal Care and Use Committee of Baylor College of Medicine. Fourteen mice (Mus musculus, 6 females, 8 males, age 2.2-4 months) expressing GCaMP6s in excitatory neurons via Slc17a7-Cre and Ai162 transgenic lines (recommended and generously shared by Hongkui Zeng at Allen Institute for Brain Science; JAX stock 023527 and 031562, respectively) were anesthetized and a 4 mm craniotomy was made over the visual cortex of the right hemisphere as described previously (Reimer et al., 2014; Froudarakis et al., 2014). Animals were allowed at least 5 days to recover before experimental scans. Mice were head-mounted above a cylindrical treadmill and two-photon calcium imaging was performed using Chameleon Ti-Sapphire laser (Coherent) tuned to 920 nm and a large field of view mesoscope (Sofroniew et al., 2016) equipped with a custom objective (excitation NA 0.6, collection NA 1.0, 21 mm focal length). Laser power after the objective was increased exponentially as a function of depth from the surface according to: 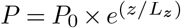, where P is the laser power used at target depth z, P0 is the power used at the surface (not exceeding 20 mW), and Lz is the depth constant (220 mm). The greatest laser output of 100 mW was used at approximately 420 mm from the surface.

The craniotomy window was leveled with regards to the objective with six degrees of freedom. Pixel-wise responses from an ROI spanning the cortical window (*>*2400 × 2400 µm, 2-5 µm/px, between 100-220 µm from surface, *>*2.47 Hz) to drifting bar stimuli were used to generate a sign map for delineating visual areas (Garrett et al., 2014). Area boundaries on the sign map were manually annotated.

For eleven out of fifteen scans (including four of the foundation cohort scans), our target imaging site was a 1200 × 1100 µm^2^ area spanning L2-L5 at the conjunction of lateral primary visual cortex (V1) and three lateral higher visual areas: anterolateral (AL), lateromedial (LM), and rostrolateral (RL). This resulted in an imaging volume that was roughly 50% V1 and 50% higher visual area. This target was chosen in order to mimic the area membership and functional property distribution in the MICrONS animal (The MICrONs Consortium et al., 2023) Each scan was performed at 6.3 Hz, collecting eight 620 × 1100 µm^2^ fields per frame at 2.5 µm/px xy resolution to tile a 1200-1220 × 1100 µm^2^ FOV at four depths (two planes per depth, 20-40 µm overlap between coplanar fields. The four imaging planes were distributed across layers with at least 45 µm spacing, with two planes in L2/3 (depths: 170-200 µm and 215-250 µm), one in L4 (300-325 mm), and one in L5 (390-420 µm).

For the remaining 4 foundation cohort scans, our target imaging site was a single plane in L2/3 (depths 210-220 µm), spanning all visual cortex visible in the cortical window (typically including V1, LM, AL, RL, PM, and AM). Each scan was performed at 6.8-6.9 Hz, collecting four 630 µm width adjacent fields (spanning 2430 µm ROI, with 90 µm total overlap). Each field was a custom height (2010-3000 µm) in order to encapsulate visual cortex within that field. Imaging was performed at 3 µm/px.

Movie of the animal’s eye and face was captured throughout the experiment. A hot mirror (Thorlabs FM02) positioned between the animal’s left eye and the stimulus monitor was used to reflect an IR image onto a camera (Genie Nano C1920M, Teledyne Dalsa) without obscuring the visual stimulus. The position of the mirror and camera were manually calibrated per session and focused on the pupil. Field of view was manually cropped for each session. The field of view contained the left eye in its entirety, and was captured at ∼20 Hz. Frame times were time stamped in the behavioral clock for alignment to the stimulus and scan frame times. Video was compressed using Labview’s MJPEG codec with quality constant of 600 and stored the frames in AVI file.

Light diffusing from the laser during scanning through the pupil was used to capture pupil diameter and eye movements. A DeepLabCut model (Mathis et al., 2018) was trained on 17 manually labeled samples from 11 animals to label each frame of the compressed eye video (intraframe only H.264 compression, CRF:17) with 8 eyelid points and 8 pupil points at cardinal and intercardinal positions. Pupil points with likelihood *>*0.9 (all 8 in 72-99% of frames per scan) were fit with the smallest enclosing circle, and the radius and center of this circle was extracted. Frames with *<* 3 pupil points with likelihood *>*0.9 (<1.2% frames per scan), or producing a circle fit with outlier *>* 5.5 standard deviations from the mean in any of the three parameters (center x, center y, radius, *<*0.2% frames per scan) were discarded (total *<*1.2% frames per scan). Gaps of *<*= 10 discarded frames were replaced by linear interpolation. Trials affected by remaining gaps were discarded (*<*18 trials per scan, *<*0.015%).

The mouse was head-restrained during imaging but could walk on a treadmill. Rostro-caudal treadmill movement was measured using a rotary optical encoder (Accu-Coder 15T-01SF-2000NV1ROC-F03-S1) with a resolution of 8000 pulses per revolution, and was recorded at ∼100 Hz in order to extract locomotion velocity. The treadmill recording was low-pass filtered with a Hamming window to remove highfrequency noise.

### Monitor positioning and calibration

Visual stimuli were presented with Psychtoolbox in MATLAB to the left eye with a 31.0 × 55.2 cm (height x width) monitor (ASUS PB258Q) with a resolution of 1080 × 1920 pixels positioned 15 cm away from the eye. When the monitor is centered on and perpendicular to the surface of the eye at the closest point, this corresponds to a visual angle of 3.8 °/cm at the nearest point and 0.7 °/cm at the most remote corner of the monitor. As the craniotomy coverslip placement during surgery and the resulting mouse positioning relative to the objective is optimized for imaging quality and stability, uncontrolled variance in animal skull position relative to the washer used for head-mounting was compensated with tailored monitor positioning on a six dimensional monitor arm. The pitch of the monitor was kept in the vertical position for all animals, while the roll was visually matched to the roll of the animal’s head beneath the headbar by the experimenter. In order to optimize the translational monitor position for centered visual cortex stimulation with respect to the imaging field of view, we used a dot stimulus with a bright background (maximum pixel intensity) and a single dark square dot (minimum pixel intensity). Randomly ordered dot locations drawn from either a 5 × 8 grid tiling the screen (20 repeats) or a 10 × 10 grid tiling a central square (approx 90 degrees width and height, 10 repeats), with each dot presentation lasting 200 ms. For five scans (four foundation cohort scans, 1 scan from Fig. 4), this dot-mapping scan targeted the V1/RL/AL/LM conjunction, and the final monitor position for each animal was chosen in order to maximize inclusion of the population receptive field peak response in cortical locations spanning the scan FOV. In the remaining scans, the procedure was the same, but the scan FOV spanned all of V1 and some adjacent higher visual areas, and thus the final monitor position for each animal was chosen in order to maximize inclusion of the population receptive field peak response in cortical locations corresponding to the extremes of the retinotopic map. In both cases, the yaw of the monitor visually matched to be perpendicular to and 15 cm from the nearest surface of the eye at that position.

A photodiode (TAOS TSL253) was sealed to the top left corner of the monitor, and the voltage was recorded at 10 KHz and timestamped with a 10 MHz behavior clock. Simultaneous measurement with a luminance meter (LS-100 Konica Minolta) perpendicular to and targeting the center of the monitor was used to generate a lookup table for linear interpolation between photodiode voltage and monitor luminance in cd/m^2^ for 16 equidistant values from 0-255, and one baseline value with the monitor unpowered.

At the beginning of each experimental session, we collected photodiode voltage for 52 full-screen pixel values from 0 to 255 for one second trials. The mean photodiode voltage for each trial was collected with an 800 ms boxcar window with 200 ms offset. The voltage was converted to luminance using previously measured relationship between photodiode voltage and luminance and the resulting luminance vs. voltage curve was fit with the function *L* = *B* + *A · P* ^*γ*^ where L is the measured luminance for pixel value P, and the median *γ* of the monitor was fit as 1.73 (range 1.58 - 1.74). All stimuli were shown without linearizing the monitor (i.e. with monitor in normal gamma mode).

During the stimulus presentation, display frame sequence information was encoded in a 3 level signal, derived from the photodiode, according to the binary encoding of the display frame (flip) number assigned in-order. This signal underwent a sine convolution, allowing for local peak detection to recover the binary signal together with its behavioral time stamps. The encoded binary signal was reconstructed for *>*96% of the flips. Each flip was time stamped by a stimulus clock (MasterClock PCIe-OSC-HSO-2 card). A linear fit was applied to the flip timestamps in the behavioral and stimulus clocks, and the parameters of that fit were used to align stimulus display frames with scanner and camera frames. The mean photodiode voltage of the sequence encoding signal at pixel values 0 and 255 was used to estimate the luminance range of the monitor during the stimulus, with minimum values of approximately 0.005 - 1 cd/m^2^ and maximum values of approximately 8.0 - 11.5 cd/m^2^.

### Scan and behavioral data preprocessing

Scan images were processed with the CAIMAN pipeline (Giovannucci et al., 2019), as described in (The MICrONs Consortium et al., 2023), to produce the spiking activity neurons at the scan rate of 6.3–6.9 Hz. The neuronal and behavioral (pupil and treadmill) activity were resampled via linear interpolation to 29.967 Hz, to match the presentation times of the stimulus video frames.

### Stimulus composition

We used dynamic libraries of natural movies and directional pink noise (“Monet”) as described in The MICrONs Consortium et al. 2023, and the static natural image library as described in Walker et al. 2019.

Dynamic Gabor filters were generated as described in Petkov and Subramanian 2007. We used a spatial envelope that had a standard deviation of approximately 16.4° in the center of the monitor. A 10-second trial consisted of 10 Gabor filters (each lasting 1 second) with randomly sampled spatial positions, directions of motion, phases, spatial and temporal frequencies.

Random dot kinematograms were generated as described in Morrone et al. 2000. The radius of the dots was approximately 2.6° in the center of the monitor. Each 10-second trial contained 5 patterns of optical flow, each lasting 2 seconds. The patterns were randomly sampled in terms of type of optical flow (translation: up/down/right/left, radial: in/out, rotation: clockwise/anticlockwise), and coherence of random dots (50%, 100%).

The stimulus compositions of the MICrONS recording sessions is described in The MICrONs Consortium et al. (2023). For all other recording session, the stimulus compositions are listed in table 1.

**Table 1.**
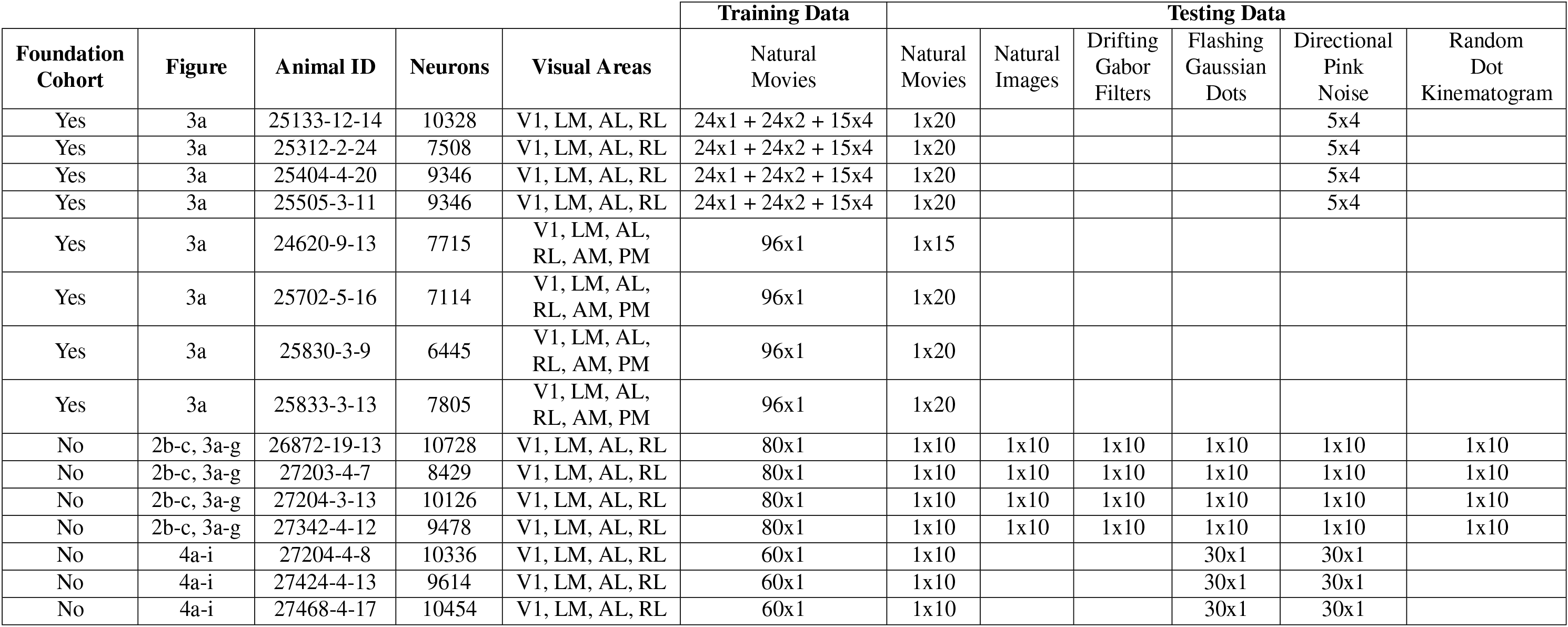
Table listing the experimental recordings, collected for either foundation core training (Foundation Cohort = Yes) or validation (Foundation Cohort = No). The animal ID, number of neurons, and areas of the visual cortex are listed for each experiment. The “Training Data” and “Testing Data” columns list the Minutes x Repeats of each type of stimulus, designated for either model training or testing.

### Neural network architecture

Our model of the visual cortex is an artificial neural network composed of four modules: perspective, behavior, core, and readout. These modules are described in the following sections.

### Perspective module

The perspective module uses ray tracing to infer the perspective or retinal activation of a mouse at discrete time points from two input variables: stimulus (movie frame) and eye position (estimated center of pupil, extracted from the eye tracking camera). To perform ray tracing, we modeled the following physical entities: 1) topography and light ray trajectories of the retina; 2) rotation of the retina; 3) position of the monitor relative to the retina; 4) intersection of the light rays of the retina and the monitor.

1) We modeled the retina as a uniform 2D grid mapped onto a 3D sphere via an azimuthal equidistant projection (Extended Data Fig. 1a). Let θ and ϕ denote the polar coordinates (radial and angular, respectively) of the 2D grid. The following mapping produces a 3D light ray for point (θ, *ϕ*) of the modeled retina:

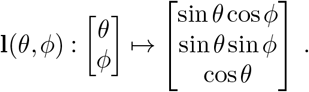

2) We used pupil tracking data to infer the rotation of the occular globe and the retina. At each time point *t*, a multilayer perceptron (MLP with 3 layers and 8 hidden units per layer) is used to map the pupil position onto the 3 ocular angles of rotation:

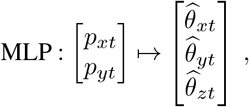

where the *p*_*xt*_, *p*_*yt*_ are the *x, y* coordinates of the pupil center in the frame of the tracking camera at time *t*, and 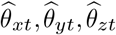 are the estimated angles of rotation of about the *x* (adduction/abduction), *y* (elevation/depression), *z* (intorsion/extorsion) axes of the occular globe at time *t*.

Let *R*_*x*_, *R*_*y*_, *R*_*z*_ ∈ ℝ^3×3^ denote rotation matrices about *x, y, z* axes. Each light ray of the retina **l**(θ, *ϕ*) is rotated by the occular angles of rotation:

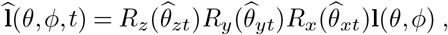

producing 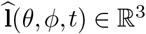, the ray of light for point (θ, *ϕ*) of the retina at time *t*, which accounts for the animal’s gaze and the rotation of the occular globe.

3) We modeled the monitor as a plane with 6 degrees of freedom: 3 for translation and 3 for rotation. Translation of the monitor plane relative to the retina is parameterized by **m**_0_ ∈ ℝ^3^. Rotation is parameterized by angles 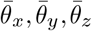:

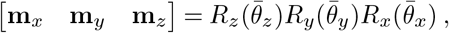

where **m**_*x*_, **m**_*y*_, **m**_*z*_ ∈ ℝ^3^ are the horizontal, vertical, and normal unit vectors of the monitor, respectively.

4) We computed the line-plane intersection between the monitor plane and 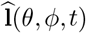, the gaze-corrected trajectory of light for point *ij* of the retina at time *t*:

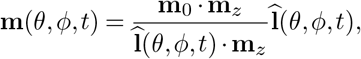

where **m**(θ, *ϕ, t*) is the point of intersection between the monitor plane and the light ray 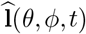. This is projected onto the monitor’s horizontal and vertical unit vectors:

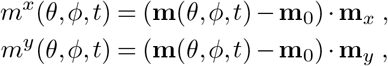

yielding *m*^*x*^(θ, *ϕ, t*) and *m*^*y*^(θ, *ϕ, t*), the horizontal and vertical displacements from the center of the monitor/stimulus (Extended Data Fig. 1b). To produce inferred activation of the retinal grid at (θ, *ϕ, t*), we performed bilinear interpolation of the stimulus at the four pixels surrounding the line-plane intersection at *m*^*x*^(θ, *ϕ, t*), *m*^*y*^(θ, *ϕ, t*).

### Modulation module

The modulation module is a small LSTM network (Hochreiter and Schmidhuber, 1997) that transforms behavioral variables, i.e., locomotion and pupil size, and previous states of the network, to produce dynamic representations of the behavioral state and arousal of the mouse.

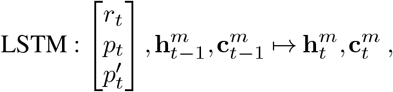

where *r* is the running/treadmill speed, *p* is the pupil diameter, *p*^′^ is the instantaneous change in pupil diameter, and **h**^*m*^, **c**^*m*^ *∈* ℝ^8^ are the “hidden” and “cell” state vectors of the modulation LSTM network.

The hidden state vector **h**^*m*^ is tiled across space to produce modulation feature maps 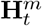:

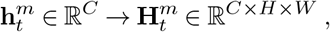

where *C, H, W* denote channel, height, and width, respectively, of the feature maps. These feature maps 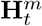 serve as the modulatory inputs into the recurrent portion of the core module at time *t*.

### Core module

The core module—comprised of feedforward and recurrent components—transforms the inputs from the perspective and modulation modules to produce feature representations of vision modulated by behavior.

First, the feedforward module transforms the visual input provided by the perspective module. For this we used DenseNet architecture (Huang et al., 2018) with 3 blocks. Each block contains 2 layers of 3D (spatiotemporal) convolutions followed by a GeLU nonlinearity ((Hendrycks and Gimpel, 2020)) and dense connections between layers. After each block, spatial pooling was performed to reduce the height and width dimensions of the feature maps. To enforce causality, we shifted the 3D convolutions along the temporal dimension, such that no inputs from future time points contributed to the output of the feedforward module.

Next, the recurrent module transforms the visual and behavioral information provided by the feedforward and modulation modules, respectively, through a group of recurrent cells. We used a convolutional LSTM (Conv-LSTM, SHI et al. 2015) as the architecture for each recurrent cell. For each cell *c*, the formulation of the Conv-LSTM is shown below:

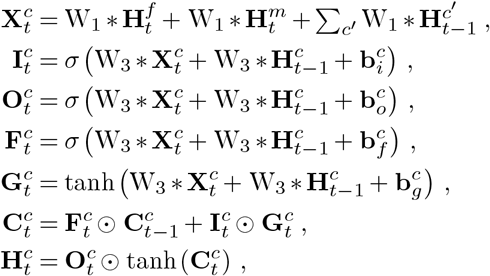

where σ denotes the sigmoid function, ⊙ denotes the Hadarmard product, and 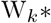 denotes a 2D spatial convolution with a *k × k* kernel. 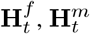 are the feedforward and modulation outputs, respectively, at time *t*, and 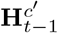 is the hidden state of an external cell *c*^*′*^ at time *t* − 1. For cell *c* at time *t*, 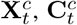 are the input, cell, and hidden states, respectively, and 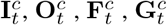 are the input, output, forget, and cell gates.

To produce the output of the core network, the hidden feature maps of the recurrent cells are concatenated along the channel dimension:

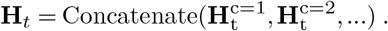

Given the recent popularity and success of transformer networks (Vaswani et al., 2023), we explored if adding the attention mechanism to our network would improve performance. We modified the Conv-Lstm architecture to incorporate the attention mechanism from the convolutional vision transformer (CvT, Wu et al. 2021). This recurrent transformer architecture, which we name CvT-Lstm, is described as follows:

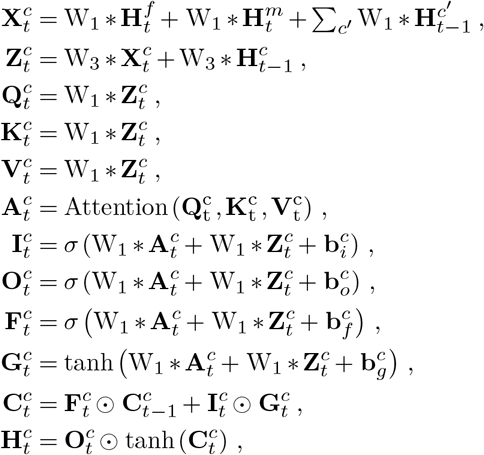

where attention is performed over query 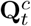, key 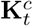, and value 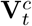 spatial tokens, which are produced by convolutions of the feature map 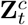. The technique of using convolutions with the attention mechanism was introduced with CvT (Wu et al., 2021), and here we extend it by incorporating it into a recurrent Lstm architecture (CvT-Lstm).

We compare the performance of Conv-Lstm vs CvT-Lstm recurrent architecture in Extended Data Fig 5. When trained on the full amount of data, Conv-Lstm performs very similarly to CvT-Lstm. However, Conv-Lstm outperforms CvT-Lstm when trained on restricted data (e.g. 4 minutes of natural movies). This was consistent for all stimulus domains that were used to test model accuracy – natural movies (Extended Data Fig 5a), natural images (b), drifting gabor filters (c), flashing gaussian dots (d), directional pink noise (e), and random dot kinematograms (e). The performance difference under data constraints may be due a better inductive bias of the Conv-Lstm. Alternatively, it could be due to a lack of optimization of the CvT-Lstm hyperparameters, and a more extensive hyperparameter search may yield better performance.

### Readout module

The readout module maps the core’s outputs onto the activity of individual neurons. For each neuron, the readout parameters are factorized into two components: spatial position and feature weights. For a neuron *n*, let **p**^*n*^ ∈ℝ^2^ denote the spatial position (*x, y*), and let **w**^*n*^ *∈ ℝ*^*C*^ denote the feature weights for that neuron, with *C* = 512 being the number channels in the core module’s output. To produce the response of that neuron *n* at time *t*, the following readout operation is performed:

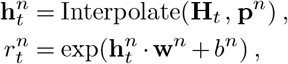

where 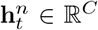 is a feature vector that is produced via bilinear interpolation of the core network’s output **H**_*t*_ *∈* ℝ^*C*×*H*×*W*^ (channels, height, width), interpolated at the spatial position **p**^*n*^. The feature vector 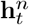 is then combined with the feature weights **w**^*n*^ and a scalar bias *b*^*n*^ to produce the response 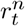 of neuron *n* at time *t*.

Due to the bilinear interpolation at a single position, each neuron only reads out from the core’s output feature maps within a 2 × 2 spatial window. While this adheres to the functional property of spatial selectivity exhibited by neurons in the visual cortex, the narrow window limits exploration of the full spatial extent of features during model training. To facilitate the spatial exploration of the core’s feature maps during training, for each neuron *n*, we sampled the readout position from a 2D Gaussian distribution: **p**^*n*^ ∼ 𝒩 (***µ***^*n*^, *Σ*^*n*^). The parameters of the distribution ***µ***^*n*^, *Σ*^*n*^ (mean, covariance) were learned via the reparameterization trick (Kingma and Welling, 2013). We observed empirically that the covariance Σ^*n*^ naturally decreased to small values by the end of training, meaning that the readout converged on a specific spatial position. After training, and for all testing purposes, we used the mean of the learned distribution ***µ***^*n*^ as the single readout position **p**^*n*^ for neuron *n*.

### Model training

The perspective, behavior, core, and readout modules were assembled together to form a model that was trained to match the recorded dynamic neuronal responses from the training dataset. Let 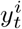 be the recorded *in vivo* response, and let 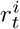 be the predicted *in silico* response of neuron *i* at time *t*. The ANN was trained to minimize the Poisson negative log likelihood loss, 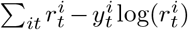, via stochastic gradient descent with Nesterov momentum (Sutskever et al., 2013). The ANN was trained for 200 epochs with a learning rate schedule that consisted of a linear warmup in the first 10 epochs, cosine decay (Loshchilov and Hutter, 2016) for 90 epochs, followed by a warm restart and cosine decay for the remaining 100 epochs. Each epoch consisted of 512 training iterations / gradient descent steps. We used a batch size of 5, and each sample of the batch consisted of 70 frames (2.33 seconds) of stimulus, neuronal, and behavioral data.

### Model hyperparameters

We used a grid search to identify architecture and training hyperparameters. Model performances for different hyperparameters were evaluated using a preliminary set of mice. After optimal hyperparameters were identified, we used the same hyperparameters to train models on a separate set of mice, from which the figures and results were produced. There was no overlap in the mice and experiments used for hyperparameter search and the mice and experiments used for the final models, results, and figures. This was done to prevent overfitting and to ensure that model performance did not depend on hyperparameters that were fit specifically for certain mice.

### Model testing

We generated model predictions of responses to stimuli that were included in the experimental recordings but excluded from model training. To evaluate the accuracy of model predictions, for each neuron we computed the correlation between the mean *in silico* and *in vivo* responses, averaged over stimulus repeats. The average *in vivo* response aims to estimate the true expected response of the neuron. However, when the *in vivo* response is highly variable and there are a limited number of repeats, this estimate becomes noisy. To account for this, we normalized the correlation by an upper bound proposed by Schoppe et al. 2016. Using 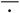 to denote average over trials/stimulus repeats, the normalized correlation *CC*_norm_ is defined as follows:

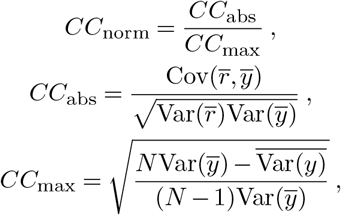

where *r* is the *in silico* response, *y* is the *in vivo* response, and *N* is the number of trials. *CC*_*abs*_ is the Pearson correlation coefficient between the average *in silico* and *in vivo* responses. *CC*_*max*_ is the upper bound of achievable performance given the the *in vivo* variability of the neuron and the number of trials.

### Parametric tuning

To estimate parametric tuning, we presented parametric stimuli to the mice and the models. Specifically, we used directional pink noise parameterized by direction/orientation and flashing Gaussian blobs parameterized by spatial location. Orientation, direction, and spatial tuning were computed from the recorded responses from the mice and the predicted responses from the models. This resulted in analogous *in vivo* and *in silico* estimates of parametric tuning for each neuron. The methods for measuring the tuning to orientation, direction, and spatial location are explained in the following sections.

### Orientation and Direction tuning

We presented 16 angles of directional pink noise, uniformly distributed between [0, 2π). Let 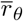 be the mean response of a neuron to the angle θ, averaged over repeated presentations of the angle. The orientation and direction selectivity indices (OSI and DSI) were computed as

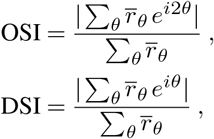

i.e., the normalized magnitude of the first and second Fourier components.

To determine the parameters for orientation and direction tuning, we used the following parametric model:

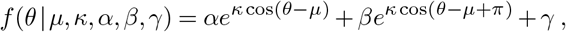

which is a mixture of two von Mises functions with amplitudes *α* and *β*, preferred directions *µ* and *µ* + π, and dispersion *κ*, plus a baseline offset of *γ*. The preferred orientation is the angle that is orthogonal to *µ* between [0, *π*], i.e., (*µ*+π*/*2) mod π. To estimate the parameters *µ, κ α, β, γ* that best fit the neuronal response, we performed least squares optimization, minimizing Σ θ (*f* (θ | *µ, κ α, β, γ*) − *r*_*θ*_)^2^.

Parameters were estimated via least square optimization for both the *in vivo* and *in silico* responses. Let 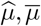 be the angles of preferred directions estimated from *in vivo, in silico* responses, respectively. The angular distances between the *in vivo* and *in silico* estimates of preferred direction (Fig. 4g) and orientation (Fig. 4e) were computed as follows:

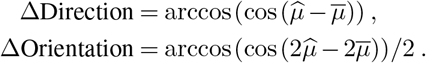

### Spatial tuning

To measure spatial tuning, we presented “on” and “off” (white and black), flashing (300 ms) Gaussian dots. The dots were isotropically shaped, with a standard deviation of approximately 8 visual degrees in the center of the monitor. The position of each dot was randomly sampled from a 17 × 29 grid tiling the height and width monitor. We observed a stronger neuronal response for “off” compared to “on”, and therefore we used only the “off” Gaussian dots to perform spatial tuning from the *in vivo* and *in silico* responses.

To measure spatial tuning, we first computed the spike triggered average (STA) of the stimulus. Let **x** ∈ ℝ^2^ denote the spatial location (height and width) in pixels. The value of the STA at location **x** was computed as follows:

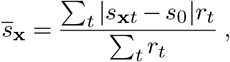

where *r*_*t*_ is the response of the neuron, *s*_**x***t*_ is the value of the stimulus at location **x** and time *t*, and *s*_0_ is the blank or gray value of the monitor.

To measure the spatial selectivity of a neuron, we computed the covariance matrix or dispersion of the STA. Again using **x** ∈ ℝ^2^ denote the spatial location (height and width) in pix-els:

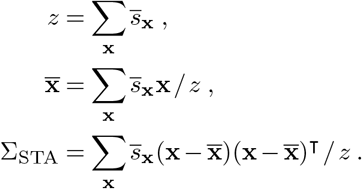

The spatial selectivity index, or strength of spatial tuning, was defined as the negative log determinant of the covariance matrix:

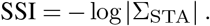

To determine the parameters of spatial tuning, we used least squares to fit the STA to the following parametric model:

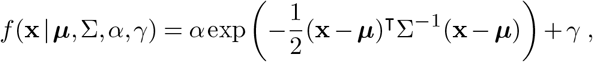

which is a 2D Gaussian component with amplitude α, mean ***µ***, and covariance Σ, plus a baseline offset of γ.

From the *in vivo* and *in silico* responses, we estimated two sets of spatial tuning parameters. Let 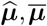 be the means (preferred spatial locations) estimated from *in vivo* and *in silico* responses. To measure the difference between the preferred locations (Fig. 4i), we computed the Euclidean distance:

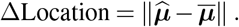

### Anatomical predictions from functional weights

To predict brain areas from readout feature weights, we used all functional units in the MICrONS data from XX scans that had readout feature weights in the model. We trained a classifier to predict brain areas from feature weights using logistic regression with nested cross validation. For each of the 10 folds, 90% of the data was used to train the model with another round of 10 fold cross validation to select the best L2 regularization weight. The best performing model was used to test on the held-out 10% of data. Finally, all of the predictions were concatenated and used to test the performance of the classifier (balanced accuracy) and generate the confusion matrix. The confusion matrix was normalized such that all rows sum to 1, thus the diagonal values represent the recall of each class. To measure the performance of V1 vs HVA’s, the balanced accuracy was measured after reclassifying the LM, RL, and AL labels in both the target and prediction as HVA.

To predict cell types, the same functional data source was used as in the brain area predictions. Cell types were obtained from CAVEclient initialized with ‘minnie65_public’ and table ‘aibs_metamodel_mtypes_v661_v2’. To associate a neuron’s functional data with its cell type, we merged the cell types to the combined manual and automatic coregistration described in (The MICrONs Consortium et al., 2023). Lastly, because each neuron could be scanned more than once, and thus could have more than one functional readout weight, we subset the data such that each neuron only had one readout weight according to its highest cc_max. Following this procedure, n=16,561 unique EM neurons remained. Out of the 20 cell classes, all excitatory neuron classes in L2-5 were chosen (except L5NP, which had comparably fewer coregistered cells), leaving 11 classes: “L2a”, “L2b”, “L2c”, “L3a”, “L3b”, “L4a”, “L4b”, “L4c”, “L5a”, “L5b”, “L5ET”. To train the classifier using readout weights to predict cell types, logistic regression was used with the same nested cross validation procedure and performance metric as described in the brain area predictions.

For testing whether readout weights contributed to cell type predictons beyond imaging depth, the 2p depth of each functional unit was obtained from a 2p structural stack (stack session 9, stack idx 19) wherein all imaging planes were registered (The MICrONs Consortium et al., 2023). This provided a common reference frame for all functional units. The two logistic regression models (depth vs depth + readout weights) were trained with all of the data, and the predicted probabilities and coefficients from the models were used to run the likelihood ratio test, where a p-value less than 0.05 was chosen as the threshold for statistical significance.

## Data availability

All MICrONS data have already been released on BossDB (https://bossdb.org/project/micronsminnie, please also see https://www.micronsexplorer.org/cortical-mm3 for details). Additional data including foundation model architecture, hyperparameters, and weights will be released upon publication.

## Code availability

All code will be released on github upon publication.

## Extended data

**Extended Data Fig. 1.**
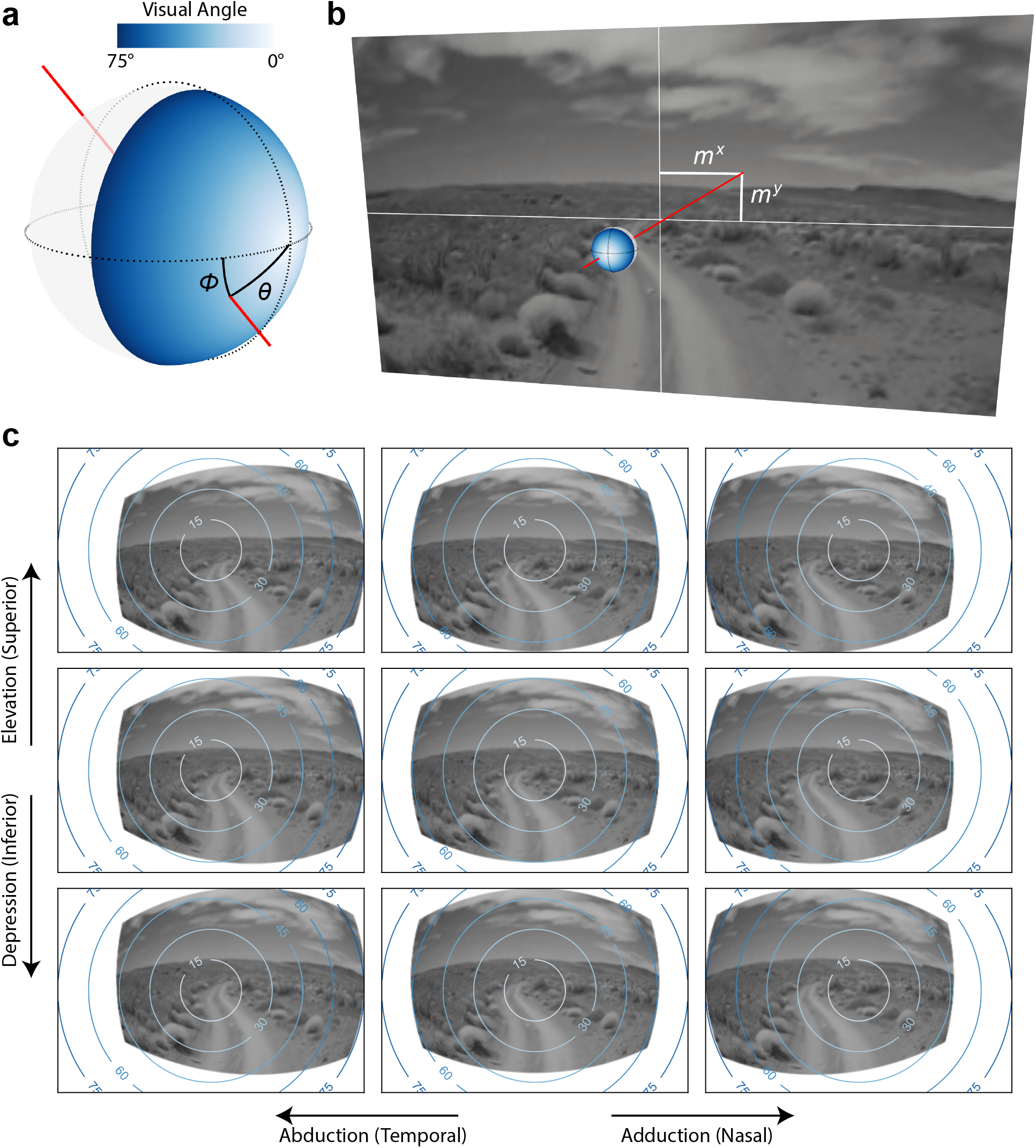
ANN perspective. Schematic of the modeled perspective the animal. **a**, The retina is modeled as points on a sphere receiving light rays that trace through the origin. An example light ray with polar angle θ and azimuthal angle ϕ is shown in red. **b**, The light ray is traced to a point *m*^*x*^,*m*^*y*^ on the monitor. Bilinear interpolation of the four pixels on the monitor surrounding *m*^*x*^,*m*^*y*^ produces the activation of a point θ, *ϕ* on the modeled retina. **c**, 9 examples of the modeled perspective from the left eye of an animal, with 3 horizontal rotations of the optical globe (abduction/adduction) × 3 vertical rotations (elevation/depression). The concentric circles indicate visual angles in degrees. (See Methods for details on the perspective network.)

**Extended Data Fig. 2.**
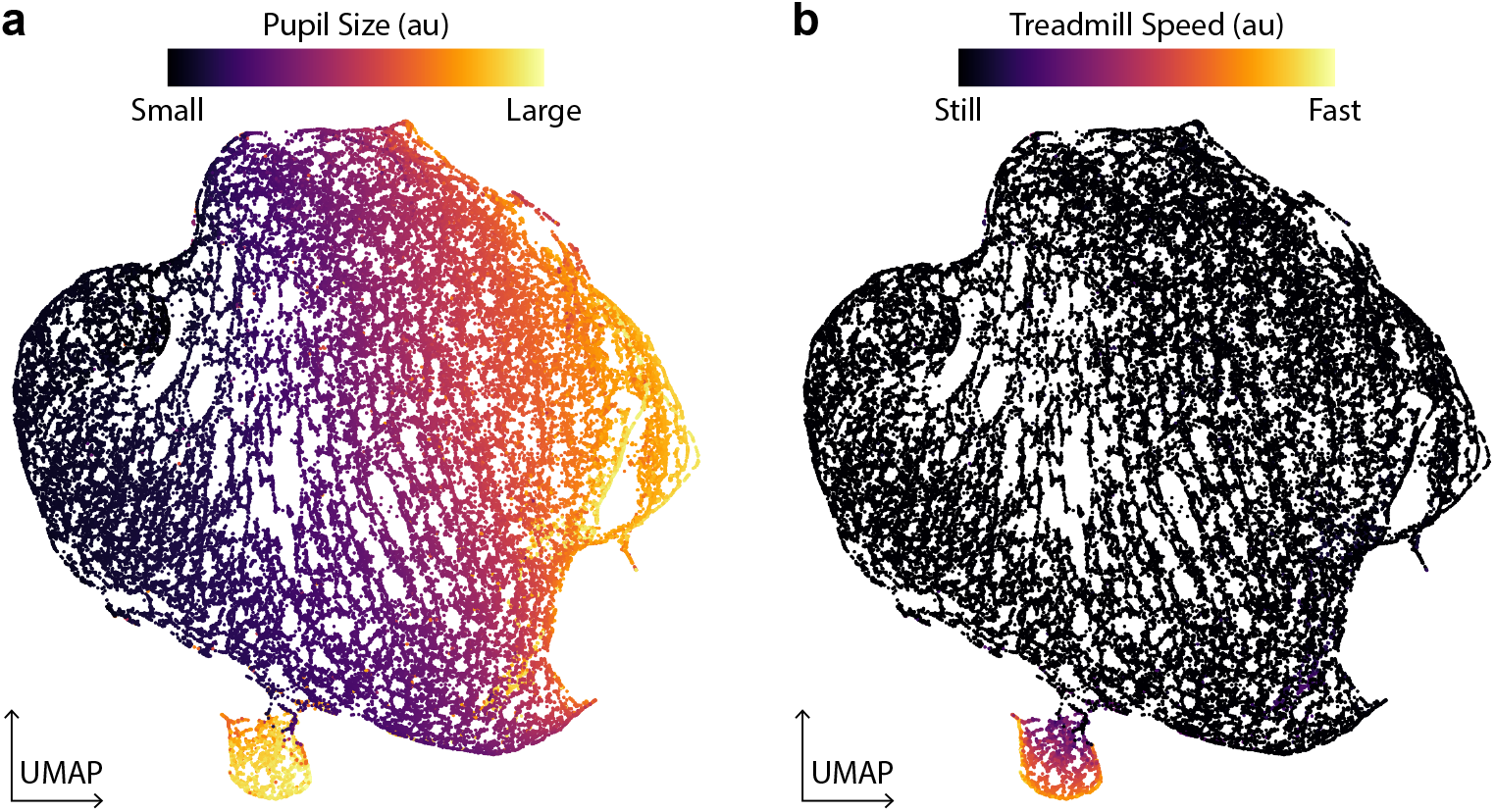
ANN modulation. Visualization of the modulation network’s output, projected onto 2 dimensions via UMAP. **a, b** show the same data from an example recording session and modulation network. Each point on the plot indicates a point in time from the recording session. The colors indicate measurements of pupil size (**a**) and treadmill speed (**b**) at the respective points in time. (See Methods for details on the modulation network.)

**Extended Data Fig. 3.**
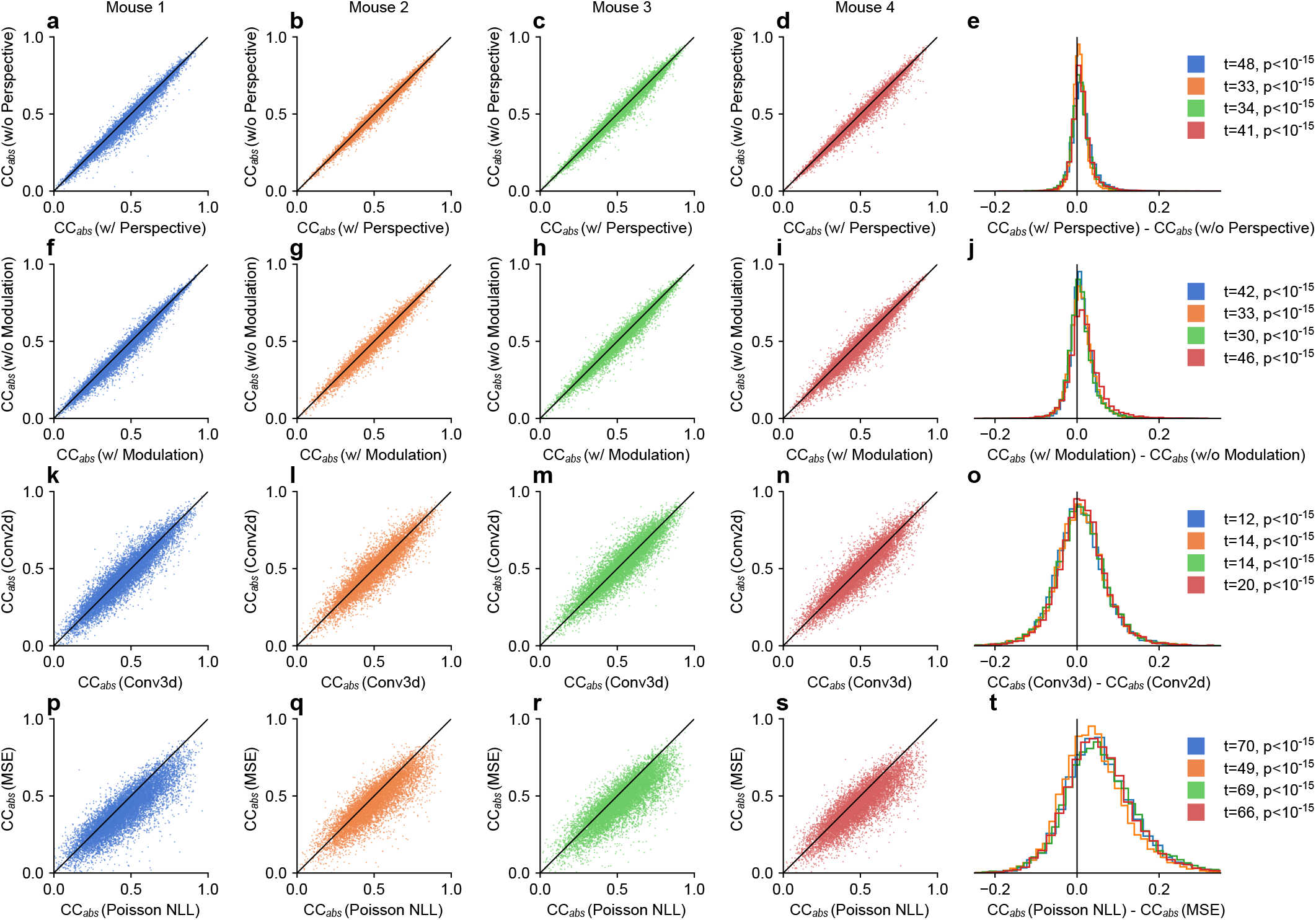
Neural network lesion studies. To determine the effect that various components of the model have on predictive accuracy, we performed lesion studies, where we altered individual components of model and evaluated the effect that the alteration had on model performance (*CC*_*abs*_). The left 4 columns (**a**-**d, f**-**i, k**-**n, p**-**s**) are scatterplots of reference vs lesioned model performance, with each column corresponding to different mouse and each point corresponding to a neuron. The right-most column (**e, j, o, t**) displays density histograms of the performance difference between the reference and the lesioned models, plotted separately for each mouse, as well as the t-statistic and p-values of paired two-sided t-tests. The first row (**a**-**e**) shows the effect of the perspective module on model performance, the second row (**f**-**j**) shows the effect of the modulation module, the third row (**k**-**o**) shows the effect of the convolution type – 2D vs 3D – of the feedforward module, and the fourth row (**p**-**t**) shows the effect of the loss function – Poisson negative log likelihood (Poission NLL) vs mean square error (MSE).

**Extended Data Fig. 4.**
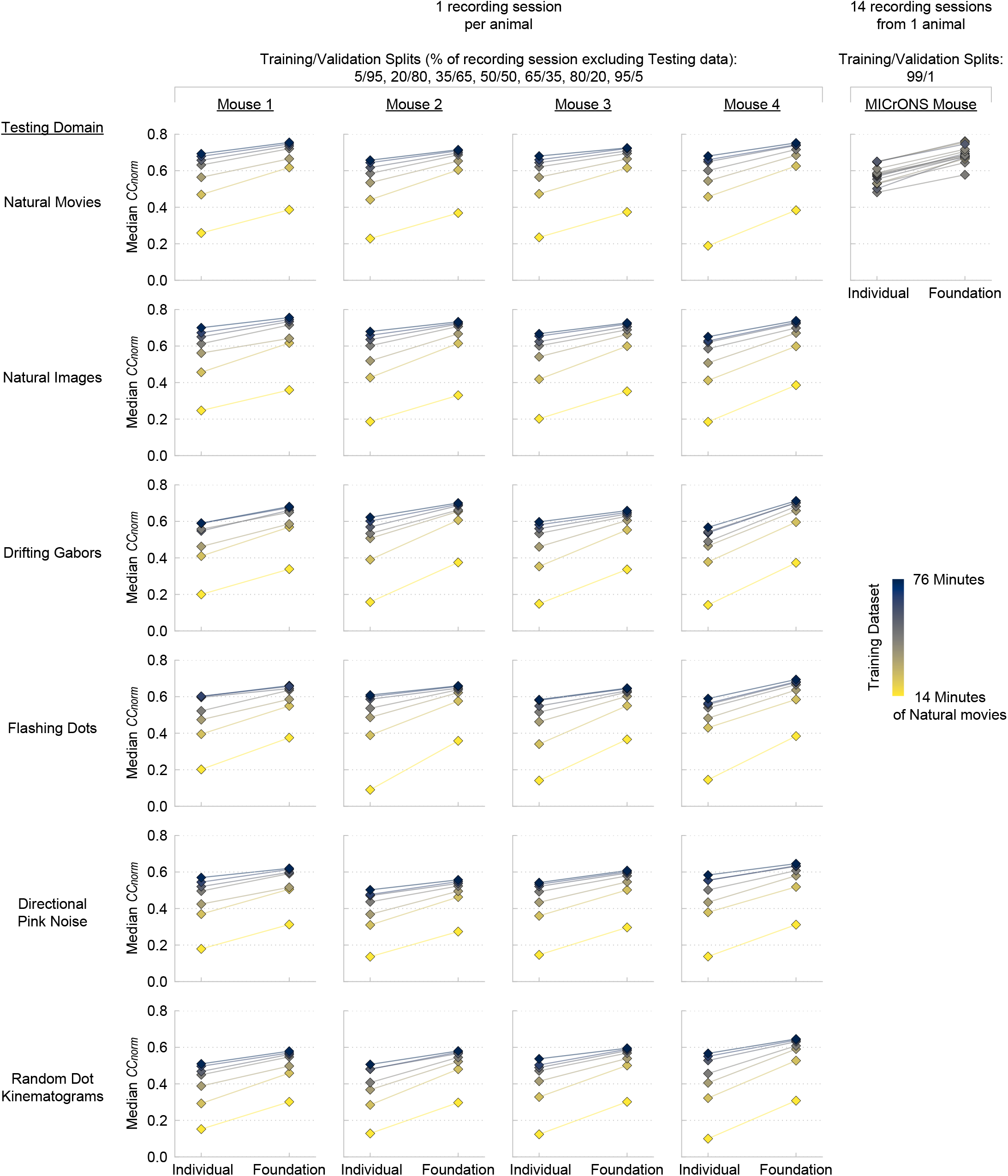
ANN performance: Individual vs. Foundation. Predictive accuracy (median *CC*_*norm*_ across neurons) of foundation models (with the foundation core) vs. individual models (with cores trained on individual recording sessions). For the 4 mice in the 4 left columns, 1 recording session was performed, and that data was partitioned into 7 training/validation splits, which were used to train separate individual/foundation models. The predictive accuracy of those models (diamonds) is reported for 6 testing stimulus domains (rows). For the MICrONS mouse, 14 recording sessions were performed, for each recording session, a model was trained using nearly all (99%) of the data available for training/validation. The MICrONS models were only tested on the natural movies, due to the lack of the other stimuli in the recording sessions. All models were trained only using natural movies.

**Extended Data Fig. 5.**
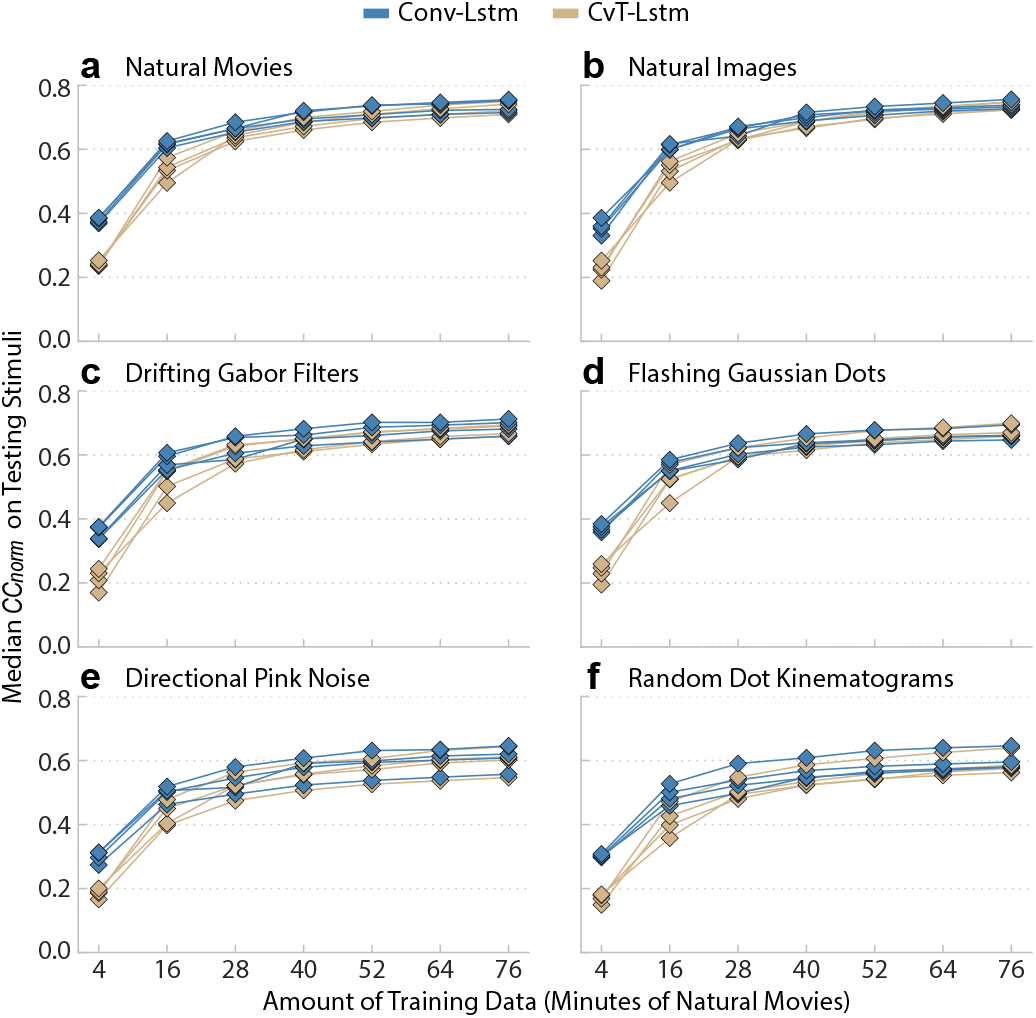
Recurrent architecture: Conv-Lstm vs. CvT-Lstm. We evaluated the performance of two different types of recurrent architectures for the core module: Conv-Lstm (blue) and CvT-Lstm (tan). For each architecture, a core was trained on 8 mice and then transferred to 4 new mice. For each of the new mice, 7 models were trained using varying amounts of natural movies, ranging from 4 to 76 minutes. The predictive accuracy (*CC*_*norm*_) of these models was evaluated on 6 different stimulus domains: natural movies (**a**), natural images (**b**), drifting gabor filter (**c**), flashing Gaussian dots (**d**), directional pink noise (**e**), random dot kinematograms (**f**). Blue diamonds indicate models with the Conv-Lstm core, and tan diamonds indicate models with the CvT-Lstm core. For each architecture, models of the same mouse are connected by lines.

**Extended Data Fig. 6.**
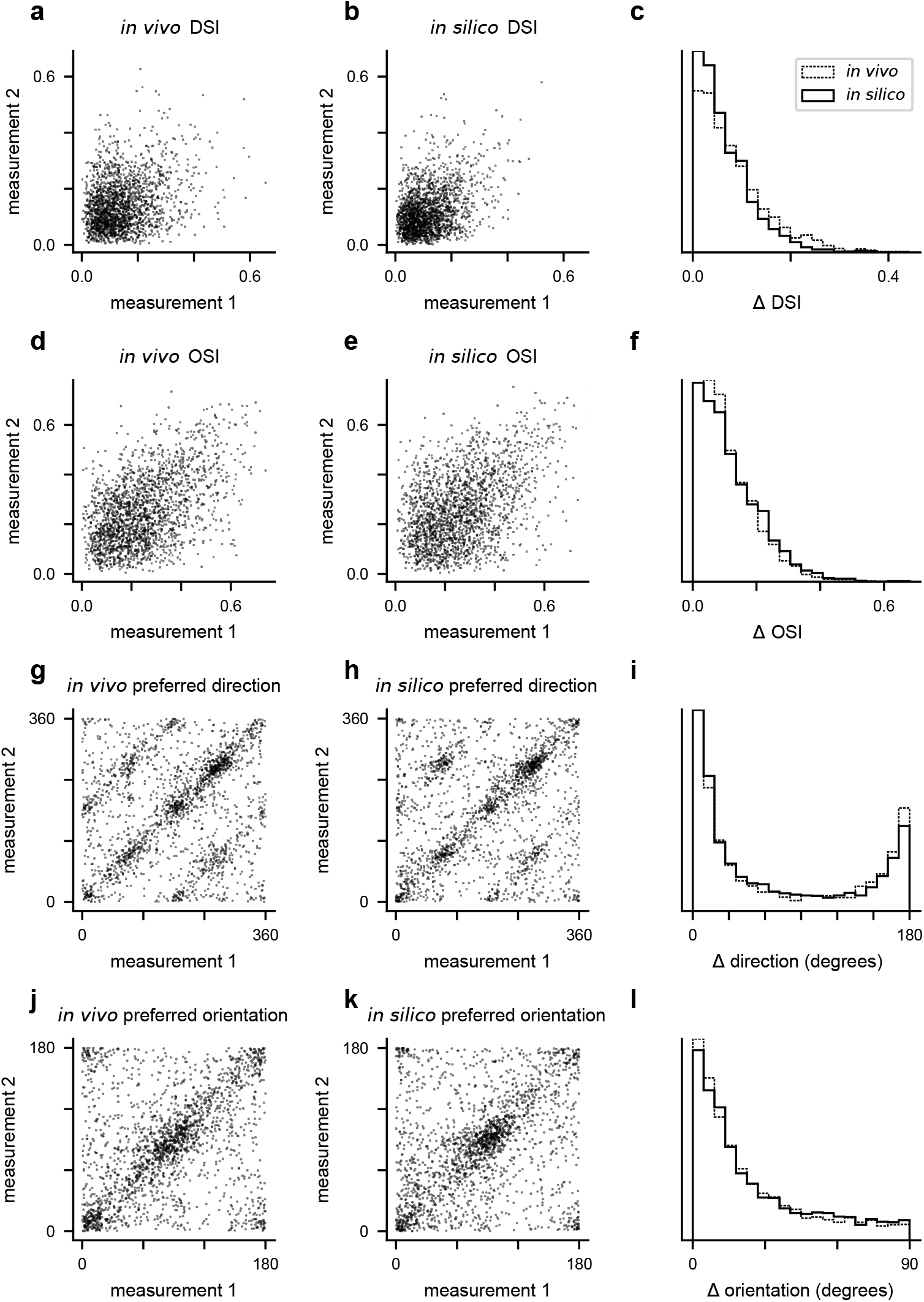
Reliability of *in vivo* and *in silico* direction and orientation tuning. **a**: Direction selectivity index (DSI) of neurons measured in two different *in vivo* experiments (i.e., recording sessions). Each point represents a single neuron measured in the two *in vivo* experiments. **b**: DSI measured in two different *in silico* experiments (i.e., model of a recording session). Each point represents a single neuron measured in the two *in silico* experiments. **c**: Distribution of the absolute differences between two measurements of DSI from *in vivo* (dashed) and *insilico* (solid) experiments. **d**-**f**: Same as **a**-**c**, but for orientation selectivity index (OSI). **g**-**i**: Same as **a**-**c**, but for preferred direction. **j**-**l**: Same as **a**-**c**, but for preferred orientation.

**Extended Data Fig. 7.**
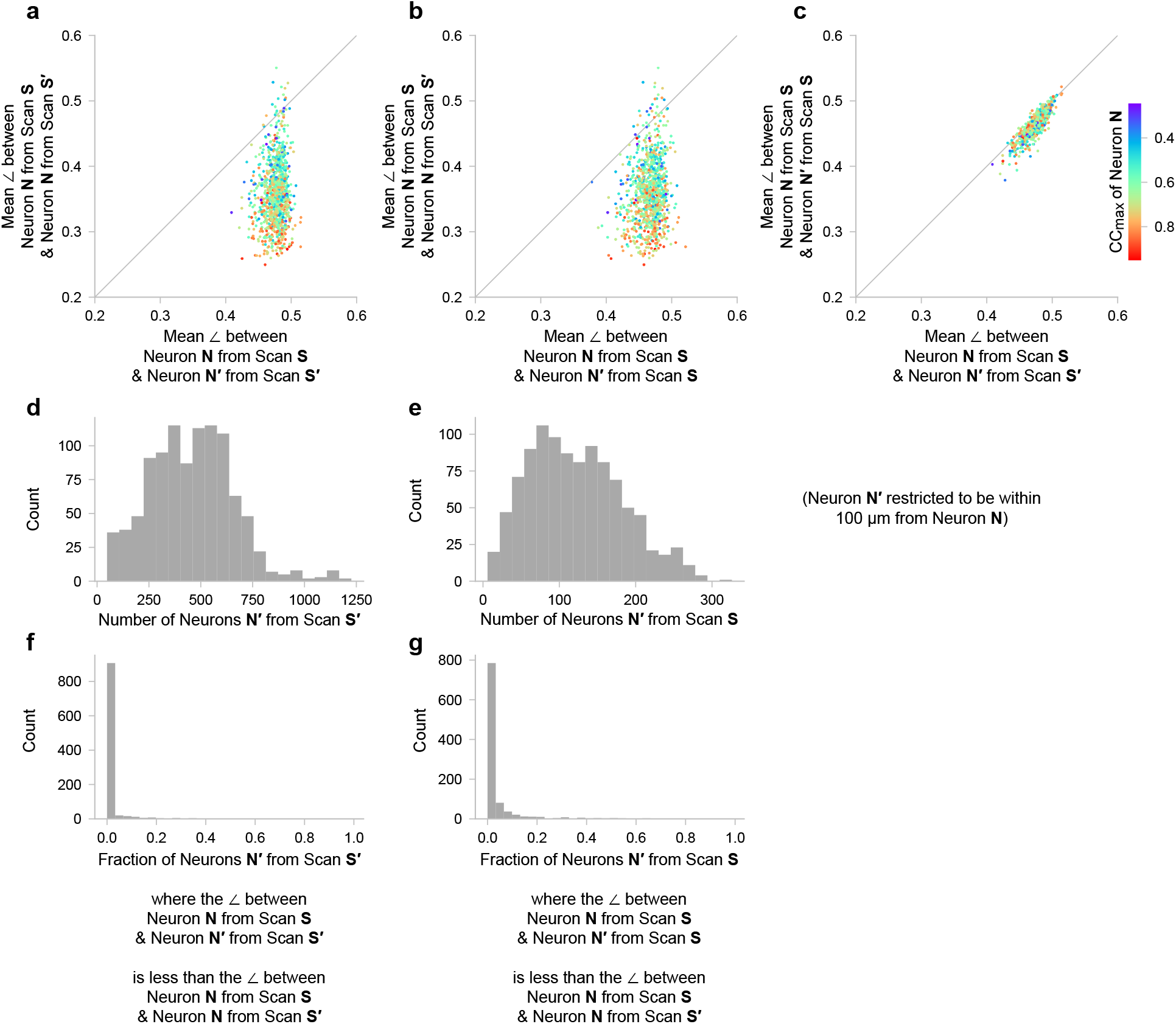
Pairwise similarities of readout feature weights of neurons from the MICrONS volume. Here we examine the similarities of readout weights of same or different neurons, from same or different scans (recording sessions). In panels **a**–**c**, the similarities of readout weights are plotted for the following groups: *same neuron* from *different scan* (y-axis of **a**), *same neuron* from *same scan* (y-axis of **b**), *different neuron* from *different scan* (x-axis of **a**, x-axis of **c**), *different neuron* from *same scan* (x-axis of **b** and y-axis of *c*). The similarity between readout weights was measured inversely via angular distance ∠ := arccos ((**x** *·* **y**)*/*(‖**x**‖**y**‖)) */π*, where **x, y** is a pair of readout weights. A similar pair of readout weights will exhibit a small ∠, and vice versa. The scatterplots **a**–**c** are colored by the *CC*_max_, which is an inverse measure of neuronal noise, i.e., the estimated maximum correlation coefficient that a model could achieve at predicting the mean response the neuron (see Methods for details). For each neuron **N**, the “different” neuron **N’** was restricted to be ≤ 100 *µ*m apart from each other in terms of soma distance, and the distribution of the number of “different” neurons is shown in **d** (from different scans) and **e** (from the same scan). **f** and **g** (corresponding to **d** and **e**, respectively) show the fraction of the nearby neurons **N’** that are more similar to **N** in terms of readout weights than **N** is to itself across different scans. **f**, For 919 out of the 1013 neurons **N**, less than 0.05 of nearby neurons **N’** from different scans had more similar readout weights. **g**, For 840 out of the 1013 neurons **N**, less than 0.05 of nearby neurons **N’** from the same scan had more similar readout weights.

## Bibliography

J. Antolík, S. B. Hofer, J. A. Bednar, and T. D. Mrsic-flogel. Model constrained by visual hierarchy improves prediction of neural responses to natural scenes. PLoS Comput. Biol., pages 1–22, 2016.

S. Bakhtiari, P. Mineault, T. Lillicrap, C. Pack, and B. Richards. The functional specialization of visual cortex emerges from training parallel pathways with self-supervised predictive learning. Advances in Neural Information Processing Systems, 34:25164–25178, 2021.

M. Bashiri, E. Walker, K.-K. Lurz, A. Jagadish, T. Muhammad, Z. Ding, Z. Ding, A. Tolias, and F. Sinz. A flow-based latent state generative model of neural population responses to natural images. Advances in Neural Information Processing Systems, 34, 2021.

Bashivan, K. Kar, and J. J. DiCarlo. Neural population control via deep image synthesis. Science (New York, N.Y.), 364(6439), 2019. ISSN 1095-9203. doi: 10.1126/science.aav9436.

Batty, J. Merel, N. Brackbill, A. Heitman, A. Sher, A. Litke, E. J. Chichilnisky, and L. Paninski. Multilayer network models of primate retinal ganglion cells. In Proceedings of the International Conference for Learning Representations (ICLR), 2017.

R. Bommasani, D. A. Hudson, E. Adeli, R. B. Altman, S. Arora, S. von Arx, M. S. Bernstein, J. Bohg, A. Bosselut, E. Brunskill, E. Brynjolfsson, S. Buch, D. Card, R. Castellon, N. S. Chatterji, A. S. Chen, K. Creel, J. Q. Davis, D. Demszky, C. Donahue, M. Doumbouya, E. Durmus, S. Ermon, J. Etchemendy, K. Ethayarajh, L. Fei-Fei, C. Finn, T. Gale, L. Gillespie, K. Goel, N. D. Goodman, S. Grossman, N. Guha, T. Hashimoto, P. Henderson, J. Hewitt, D. E. Ho, J. Hong, K. Hsu, J. Huang, T. Icard, S. Jain, D. Jurafsky, P. Kalluri, S. Karamcheti, G. Keeling, F. Khani, O. Khattab, P. W. Koh, M. S. Krass, R. Krishna, R. Kuditipudi, and et al. On the opportunities and risks of foundation models. CoRR, abs/2108.07258, 2021. URL https://arxiv.org/abs/2108.07258.

K. H. Britten, M. N. Shadlen, W. T. Newsome, and J. A. Movshon. The analysis of visual motion: a comparison of neuronal and psychophysical performance. Journal of Neuroscience, 12(12): 4745–4765, 1992.

T. B. Brown, B. Mann, N. Ryder, M. Subbiah, J. Kaplan, P. Dhariwal, A. Neelakantan, P. Shyam, G. Sastry, A. Askell, S. Agarwal, A. Herbert-Voss, G. Krueger, T. Henighan, R. Child, A. Ramesh, D. M. Ziegler, J. Wu, C. Winter, C. Hesse, M. Chen, E. Sigler, M. Litwin, S. Gray, B. Chess, J. Clark, C. Berner, S. McCandlish, A. Radford, I. Sutskever, and D. Amodei. Language models are few-shot learners. CoRR, abs/2005.14165, 2020. URL https://arxiv.org/abs/2005.14165.

F. Burg, S. A. Cadena, G. H. Denfield, E. Y. Walker, A. S. Tolias, M. Bethge, and A. S. Ecker. Learning divisive normalization in primary visual cortex. PLOS Computational Biology, 17(6): e1009028, July 2021. ISSN 1553-7358. doi: 10.1371/journal.pcbi.1009028.

S. A. Cadena, G. H. Denfield, E. Y. Walker, L. A. Gatys, A. S. Tolias, M. Bethge, and A. S. Ecker. Deep convolutional models improve predictions of macaque v1 responses to natural images. PLOS Computational Biology, 15(4):e1006897, Apr. 2019. doi: 10.1371/journal.pcbi.1006897.

C. F. Cadieu, H. Hong, D. L. K. Yamins, N. Pinto, D. Ardila, E. A. Solomon, N. J. Majaj, and J. J. DiCarlo. Deep neural networks rival the representation of primate IT cortex for core visual object recognition. PLoS Comput. Biol., 10(12):e1003963, 2014.

E. Christensen and J. Zylberberg. Models of primate ventral stream that categorize and visualize images. bioRxiv, pages 2020–02, 2020.

B. Cowley and J. Pillow. High-contrast “gaudy” images improve the training of deep neural network models of visual cortex. In H. Larochelle, M. Ranzato, R. Hadsell, M. Balcan, and H. Lin, editors, Advances in Neural Information Processing Systems 33, pages 21591–21603. Curran Associates, Inc., 2020.

Z. Ding, P. G. Fahey, S. Papadopoulos, E. Y. Wang, B. Celii, C. Papadopoulos, A. B. Kunin, A. Chang, J. Fu, Z. Ding, S. Patel, K. Ponder, T. Muhammad, J. A. Bae, A. L. Bodor, D. Brittain, J. Buchanan, D. J. Bumbarger, M. A. Castro, E. Cobos, S. Dorkenwald, L. Elabbady, A. Halageri, Z. Jia, C. Jordan, D. Kapner, N. Kemnitz, S. Kinn, K. Lee, K. Li, R. Lu, T. Macrina, G. Mahalingam, E. Mitchell, S. S. Mondal, S. Mu, B. Nehoran, S. Popovych, C. M. Schneider-Mizell, W. Silversmith, M. Takeno, R. Torres, N. L. Turner, W. Wong, J. Wu, W. Yin, S.-c. Yu, E. Froudarakis, F. Sinz, H. S. Seung, F. Collman, N. M. da Costa, R. C. Reid, E. Y. Walker, X. Pitkow, J. Reimer, and A. S. Tolias. Functional connectomics reveals general wiring rule in mouse visual cortex, 2023a. URL 10.1101/2023.03.13.531369.

Z. Ding, D. T. Tran, K. Ponder, E. Cobos, Z. Ding, P. G. Fahey, E. Wang, T. Muhammad, J. Fu, S. A. Cadena, S. Papadopoulos, S. Patel, K. Franke, J. Reimer, F. H. Sinz, A. S. Ecker, X. Pitkow, and A. S. Tolias. Bipartite invariance in mouse primary visual cortex, 2023b. URL 10.1101/2023.03.15.532836.

A. S. Ecker, F. H. Sinz, E. Froudarakis, P. G. Fahey, S. A. Cadena, E. Y. Walker, E. Cobos, J. Reimer, A. S. Tolias, and M. Bethge. A rotation-equivariant convolutional neural network model of primary visual cortex, 2018. URL https://arxiv.org/abs/1809.10504.

K. Franke, K. F. Willeke, K. Ponder, M. Galdamez, N. Zhou, T. Muhammad, S. Patel, E. Froudarakis, J. Reimer, F. H. Sinz, and A. S. Tolias. State-dependent pupil dilation rapidly shifts visual feature selectivity. 610(7930):128–134, 2022. ISSN 0028-0836, 1476-4687. doi: 10.1038/s41586-022-05270-3.

E. Froudarakis, P. Berens, A. S. Ecker, R. J. Cotton, F. H. Sinz, D. Yatsenko, P. Saggau, M. Bethge, and A. S. Tolias. Population code in mouse V1 facilitates readout of natural scenes through increased sparseness. Nat. Neurosci., 17(6):851–857, June 2014.

J. Fu, S. Shrinivasan, K. Ponder, T. Muhammad, Z. Ding, E. Wang, Z. Ding, D. T. Tran, P. G. Fahey, S. Papadopoulos, S. Patel, J. Reimer, A. S. Ecker, X. Pitkow, R. M. Haefner, F. H. Sinz, K. Franke, and A. S. Tolias. Pattern completion and disruption characterize contextual modulation in mouse visual cortex, 2023. URL 10.1101/2023.03.13.532473.

M. E. Garrett, I. Nauhaus, J. H. Marshel, and E. M. Callaway. Topography and areal organization of mouse visual cortex. J. Neurosci., 34(37):12587–12600, Sept. 2014.

A. Giovannucci, J. Friedrich, P. Gunn, J. Kalfon, B. L. Brown, S. A. Koay, J. Taxidis, F. Najafi, J. L. Gauthier, P. Zhou, B. S. Khakh, D. W. Tank, D. B. Chklovskii, and E. A. Pnevmatikakis. CaImAn: An open source tool for scalable calcium imaging data analysis. Elife, 8:e38173, 2019.

P. M. Goltstein, S. Reinert, T. Bonhoeffer, and M. Hübener. Mouse visual cortex areas represent perceptual and semantic features of learned visual categories. 2021. ISSN 1097-6256, 1546-1726. doi: 10.1038/s41593-021-00914-5.

D. Hendrycks and T. G. Dietterich. Benchmarking neural network robustness to common corruptions and perturbations. CoRR, abs/1903.12261, 2019. URL http://arxiv.org/abs/1903.12261.

D. Hendrycks and K. Gimpel. Gaussian error linear units (GELUs), 2020. URL http://arxiv.org/abs/1606.08415.

S. Hochreiter and J. Schmidhuber. Long short-term memory. 9(8):1735–1780, 1997. ISSN 0899-7667, 1530-888X. doi: 10.1162/neco.1997.9.8.1735. URL https://direct.mit.edu/neco/article/9/8/1735-1780/6109.

L. Höfling, K. P. Szatko, C. Behrens, Y. Qiu, D. A. Klindt, Z. Jessen, G. W. Schwartz, M. Bethge, P. Berens, K. Franke, A. S. Ecker, and T. Euler. A chromatic feature detector in the retina signals visual context changes. Dec. 2022.

G. Huang, Z. Liu, L. van der Maaten, and K. Q. Weinberger. Densely connected convolutional networks, 2018. URL http://arxiv.org/abs/1608.06993.

W. F. Kindel, E. D. Christensen, and J. Zylberberg. Using deep learning to reveal the neural code for images in primary visual cortex. 2017.

D. P. Kingma and M. Welling. Auto-encoding variational bayes, 2013. URL https://arxiv.org/abs/1312.6114.

D. A. Klindt, A. S. Ecker, T. Euler, and M. Bethge. Neural system identification for large populations separating “what” and “where”. In Advances in Neural Information Processing Systems, pages 4–6, 2017.

T. H. Kung, M. Cheatham, A. Medenilla, C. Sillos, L. De Leon, C. Elepaño, M. Madriaga, R. Aggabao, G. Diaz-Candido, J. Maningo, and V. Tseng. Performance of ChatGPT on USMLE: Potential for AI-assisted medical education using large language models. 2(2):e0000198, 2023. ISSN 2767-3170. doi: 10.1371/journal.pdig.0000198.

M. J. Lindstrom and D. M. Bates. Newton-raphson and EM algorithms for linear mixed-effects models for repeated-measures data. 83(404):1014, 1988. ISSN 01621459. doi: 10.2307/2290128.

I. Loshchilov and F. Hutter. SGDR: stochastic gradient descent with restarts. CoRR, abs/1608.03983, 2016. URL http://arxiv.org/abs/1608.03983.

K.-K. Lurz, M. Bashiri, K. Willeke, A. K. Jagadish, E. Wang, E. Y. Walker, S. A. Cadena, T. Muhammad, E. Cobos, A. S. Tolias, A. S. Ecker, and F. H. Sinz. Generalization in data-driven models of primary visual cortex. In Proceedings of the International Conference for Learning Representations (ICLR), page 2020.10.05.326256, Oct. 2020.

K.-K. Lurz, M. Bashiri, K. Willeke, A. Jagadish, E. Wang, E. Y. Walker, S. A. Cadena, T. Muhammad, E. Cobos, A. S. Tolias, A. S. Ecker, and F. H. Sinz. Generalization in data-driven models of primary visual cortex. In International Conference on Learning Representations, 2021.

J. H. Marshel, Y. S. Kim, T. A. Machado, S. Quirin, B. Benson, J. Kadmon, C. Raja, A. Chibukhchyan, C. Ramakrishnan, M. Inoue, et al. Cortical layer–specific critical dynamics triggering perception. Science, 365(6453):eaaw5202, 2019.

A. Mathis, P. Mamidanna, K. M. Cury, T. Abe, V. N. Murthy, M. W. Mathis, and M. Bethge. DeepLabCut: markerless pose estimation of user-defined body parts with deep learning. Nat. Neurosci., 21(9):1281–1289, Aug. 2018.

L. T. McIntosh, N. Maheswaranathan, A. Nayebi, S. Ganguli, and S. A. Baccus. Deep learning models of the retinal response to natural scenes. Adv. Neural Inf. Process. Syst., 29(Nips): 1369–1377, 2016.

M. C. Morrone, M. Tosetti, D. Montanaro, A. Fiorentini, G. Cioni, and D. C. Burr. A cortical area that responds specifically to optic flow, revealed by fMRI. Nature Neuroscience, 3(12):1322–1328, 2000. ISSN 1097-6256, 1546-1726. doi: 10.1038/81860.

A. Nayebi, N. C. Kong, C. Zhuang, J. L. Gardner, A. M. Norcia, and D. L. Yamins. Shallow unsupervised models best predict neural responses in mouse visual cortex. bioRxiv, pages 2021–06, 2021.

N. Petkov and E. Subramanian. Motion detection, noise reduction, texture suppression and contour enhancement by spatiotemporal gabor filters with surround inhibition. Biological Cybernetics, 97(5-6):423–439, 2007. doi: 10.1007/s00422-007-0182-0.

P. Pierzchlewicz, K. Willeke, A. Nix, P. Elumalai, K. Restivo, T. Shinn, C. Nealley, G. Rodriguez, S. Patel, K. Franke, et al. Energy guided diffusion for generating neurally exciting images. Advances in Neural Information Processing Systems, 36, 2024.

C. R. Ponce, W. Xiao, P. F. Schade, T. S. Hartmann, G. Kreiman, and M. S. Livingstone. Evolving images for visual neurons using a deep generative network reveals coding principles and neuronal preferences. Cell, 177(4):999–1009.e10, 2019.

A. Radford, J. W. Kim, C. Hallacy, A. Ramesh, G. Goh, S. Agarwal, G. Sastry, A. Askell, P. Mishkin, J. Clark, G. Krueger, and I. Sutskever. Learning transferable visual models from natural language supervision. CoRR, abs/2103.00020, 2021. URL https://arxiv.org/abs/2103.00020.

J. Reimer, E. Froudarakis, C. R. Cadwell, D. Yatsenko, G. H. Denfield, and A. S. Tolias. Pupil fluctuations track fast switching of cortical states during quiet wakefulness. Neuron, 84(2): 355–362, Oct. 2014.

C. D. Salzman, K. H. Britten, and W. T. Newsome. Cortical microstimulation influences perceptual judgements of motion direction. Nature, 346(6280):174–177, 1990.

C. M. Schneider-Mizell, A. Bodor, D. Brittain, J. Buchanan, D. J. Bumbarger, L. Elabbady, D. Kapner, S. Kinn, G. Mahalingam, S. Seshamani, et al. Cell-type-specific inhibitory circuitry from a connectomic census of mouse visual cortex. bioRxiv, 2023.

O. Schoppe, N. S. Harper, B. D. B. Willmore, A. J. King, and J. W. H. Schnupp. Measuring the performance of neural models. Frontiers in Computational Neuroscience, 10, Feb. 2016. doi: 10.3389/fncom.2016.00010.

X. Shi, Z. Chen, H. Wang, D.-Y. Yeung, W.-k. Wong, and W.-c. Woo. Convolutional lstm network: A machine learning approach for precipitation nowcasting. In C. Cortes, N. Lawrence, D. Lee, M. Sugiyama, and R. Garnett, editors, Advances in Neural Information Processing Systems, volume 28. Curran Associates, Inc., 2015.

J. H. Siegle, X. Jia, S. Durand, S. Gale, C. Bennett, N. Graddis, G. Heller, T. K. Ramirez, H. Choi, J. A. Luviano, P. A. Groblewski, R. Ahmed, A. Arkhipov, A. Bernard, Y. N. Billeh, D. Brown, M. A. Buice, N. Cain, S. Caldejon, L. Casal, A. Cho, M. Chvilicek, T. C. Cox, K. Dai, D. J. Denman, S. E. J. de Vries, R. Dietzman, L. Esposito, C. Farrell, D. Feng, J. Galbraith, M. Garrett, E. C. Gelfand, N. Hancock, J. A. Harris, R. Howard, B. Hu, R. Hytnen, R. Iyer, E. Jessett, K. Johnson, I. Kato, J. Kiggins, S. Lambert, J. Lecoq, P. Ledochowitsch, J. H. Lee, A. Leon, Y. Li, E. Liang, F. Long, K. Mace, J. Melchior, D. Millman, T. Mollenkopf, C. Nayan, L. Ng, K. Ngo, T. Nguyen, P. R. Nicovich, K. North, G. K. Ocker, D. Ollerenshaw, M. Oliver, M. Pachitariu, J. Perkins, M. Reding, D. Reid, M. Robertson, K. Ronellenfitch, S. Seid, C. Slaughterbeck, M. Stoecklin, D. Sullivan, B. Sutton, J. Swapp, C. Thompson, K. Turner, W. Wakeman, J. D. Whitesell, D. Williams, A. Williford, R. Young, H. Zeng, S. Naylor, J. W. Phillips, R. C. Reid, S. Mihalas, S. R. Olsen, and C. Koch. Survey of spiking in the mouse visual system reveals functional hierarchy. 2021. ISSN 0028-0836, 1476-4687. doi: 10.1038/s41586-020-03171-x.

F. Sinz, A. S. Ecker, P. Fahey, E. Walker, E. Cobos, E. Froudarakis, D. Yatsenko, X. Pitkow, J. Reimer, and A. Tolias. Stimulus domain transfer in recurrent models for large scale cortical population prediction on video. In Advances in Neural Information Processing Systems 31. 2018.

N. J. Sofroniew, D. Flickinger, J. King, and K. Svoboda. A large field of view two-photon mesoscope with subcellular resolution for in vivo imaging. Elife, 5:e14472, June 2016.

I. Sutskever, J. Martens, G. Dahl, and G. Hinton. On the importance of initialization and momentum in deep learning. In S. Dasgupta and D. McAllester, editors, Proceedings of the 30th International Conference on Machine Learning, volume 28 of Proceedings of Machine Learning Research, pages 1139–1147, Atlanta, Georgia, USA, 17–19 Jun 2013. PMLR.

The MICrONs Consortium, J. A. Bae, M. Baptiste, C. A. Bishop, A. L. Bodor, D. Brittain, J. Buchanan, D. J. Bumbarger, M. A. Castro, B. Celii, E. Cobos, F. Collman, N. M. da Costa, S. Dorkenwald, L. Elabbady, P. G. Fahey, T. Fliss, E. Froudarakis, J. Gager, C. Gamlin, W. R. Gray-Roncal, A. Halageri, J. Hebditch, Z. Jia, C. Jordan, E. Joyce, J. Joyce, D. Kapner, N. Kemnitz, S. Kinn, L. M. Kitchell, S. Koolman, K. Kuehner, K. Lee, K. Li, R. Lu, T. Macrina, G. Mahalingam, J. Matelsky, S. McReynolds, E. Miranda, E. Mitchell, S. S. Mondal, M. Moore, S. Mu, T. Muhammad, B. Nehoran, O. Ogedengbe, C. Papadopoulos, S. Papadopoulos, S. Patel, X. Pitkow, S. Popovych, A. Ramos, R. C. Reid, J. Reimer, P. K. Rivlin, C. M. Schneider-Mizell, V. Rose, H. S. Seung, B. Silverman, W. Silversmith, F. H. Sinz, C. L. Smith, A. Sterling, S. Suckow, M. Takeno, Z. H. Tan, A. S. Tolias, R. Torres, N. L. Turner, E. Y. Walker, T. Wang, A. Wanner, B. A. Wester, G. Williams, S. Williams, K. Willie, R. Willie, W. Wong, J. Wu, D. R. Xenes, C. Xu, R. Yang, D. Yatsenko, F. Ye, W. Yin, R. Young, S.-c. Yu, and C. Zhang. Functional connectomics spanning multiple areas of mouse visual cortex, 2023. URL 10.1101/2021.07.28.454025.

I. Ustyuzhaninov, M. F. Burg, S. A. Cadena, J. Fu, T. Muhammad, K. Ponder, E. Froudarakis, Z. Ding, M. Bethge, A. S. Tolias, and A. S. Ecker. Digital twin reveals combinatorial code of non-linear computations in the mouse primary visual cortex. Feb. 2022. doi: 10.1101/2022.02.10.479884. URL. https://doi.org/10.1101/2022.02.10.479884

A. Vaswani, N. Shazeer, N. Parmar, J. Uszkoreit, L. Jones, A. N. Gomez, L. Kaiser, and I. Polosukhin. Attention is all you need, 2023. URL http://arxiv.org/abs/1706.03762.

M. Vystrčilová, S. Sridhar, M. F. Burg, T. Gollisch, and A. S. Ecker. Convolutional neural network models of the primate retina reveal adaptation to natural stimulus statistics. bioRxiv, 2024. doi: 10.1101/2024.03.06.583740. URL https://www.biorxiv.org/content/early/2024/03/09/2024.03.06.583740.

E. Y. Walker, F. H. Sinz, E. Cobos, T. Muhammad, E. Froudarakis, P. G. Fahey, A. S. Ecker, J. Reimer, X. Pitkow, and A. S. Tolias. Inception loops discover what excites neurons most using deep predictive models. Nat. Neurosci., 22(12):2060–2065, Dec. 2019.

M. A. Weis, S. Papadopoulos, L. Hansel, T. Lüddecke, B. Celii, P. G. Fahey, E. Y. Wang, J. A. Bae, A. L. Bodor, D. Brittain, J. Buchanan, D. J. Bumbarger, M. A. Castro, F. Collman, N. M. Da Costa, S. Dorkenwald, L. Elabbady, A. Halageri, Z. Jia, C. Jordan, D. Kapner, N. Kemnitz, S. Kinn, K. Lee, K. Li, R. Lu, T. Macrina, G. Mahalingam, E. Mitchell, S. S. Mondal, S. Mu, B. Nehoran, S. Popovych, R. C. Reid, C. M. Schneider-Mizell, H. S. Seung, W. Silversmith, M. Takeno, R. Torres, N. L. Turner, W. Wong, J. Wu, W. Yin, S.-c. Yu, J. Reimer, P. Berens, A. S. Tolias, and A. S. Ecker. An unsupervised map of excitatory neurons’ dendritic morphology in the mouse visual cortex, 2022.

K. F. Willeke, P. G. Fahey, M. Bashiri, L. Hansel, C. Blessing, K.-K. Lurz, M. F. Burg, S. A. Cadena, Z. Ding, K. Ponder, T. Muhammad, S. S. Patel, K. Deng, Y. Guan, Y. Zhu, K. Xiao, X. Han, S. Azeglio, U. Ferrari, P. Neri, O. Marre, A. Hoffmann, K. Fedyanin, K. Vishniakov, M. Panov, S. Prakash, K. Naik, K. Narayanappa, A. S. Ecker, A. S. Tolias, and F. H. Sinz. Retrospective on the sensorium 2022 competition. In M. Ciccone, G. Stolovitzky, and J. Albrecht, editors, Proceedings of the NeurIPS 2022 Competitions Track, volume 220 of Proceedings of Machine Learning Research, pages 314–333. PMLR, 28 Nov–09 Dec 2022. URL https://proceedings.mlr.press/v220/willeke23a.html.

H. Wu, B. Xiao, N. Codella, M. Liu, X. Dai, L. Yuan, and L. Zhang. CvT: Introducing convolutions to vision transformers, 2021. URL http://arxiv.org/abs/2103.15808.

D. L. K. Yamins, H. Hong, C. F. Cadieu, E. A. Solomon, D. Seibert, and J. J. DiCarlo. Performance-optimized hierarchical models predict neural responses in higher visual cortex. Proceedings of the National Academy of Sciences, May 2014a.

D. L. K. Yamins, H. Hong, C. F. Cadieu, E. A. Solomon, D. Seibert, and J. J. DiCarlo. Performance-optimized hierarchical models predict neural responses in higher visual cortex. 111(23):8619–8624, 2014b. ISSN 0027-8424, 1091-6490. doi: 10.1073/pnas.1403112111. URL https://pnas.org/doi/full/10.1073/pnas.1403112111.

J. Zhuang, L. Ng, D. Williams, M. Valley, Y. Li, M. Garrett, and J. Waters. An extended retinotopic map of mouse cortex. eLife, 6:e18372, jan 2017. ISSN 2050-084X.

